# ZFP281 coordinates DNMT3 and TET1 for transcriptional and epigenetic control in pluripotent state transitions

**DOI:** 10.1101/2023.03.24.534143

**Authors:** Xin Huang, Sophie Balmer, Cong Lyu, Yunlong Xiang, Vikas Malik, Hailin Wang, Yu Zhang, Wei Xie, Anna-Katerina Hadjantonakis, Hongwei Zhou, Jianlong Wang

## Abstract

The progression from naive through formative to primed *in vitro* pluripotent stem cell states recapitulates the development of the epiblast *in vivo* during the peri-implantation period of mammalian development. Activation of the *de novo* DNA methyltransferases and reorganization of transcriptional and epigenetic landscapes are key events occurring during these pluripotent state transitions. However, the upstream regulators that coordinate these events are relatively underexplored. Here, using *Zfp281* knockout mouse and degron knock-in cell models, we uncover the direct transcriptional activation of *Dnmt3a/3b* by ZFP281 in pluripotent stem cells. Chromatin co-occupancy of ZFP281 and DNA hydroxylase TET1, dependent on the formation of R loops in ZFP281-targeted gene promoters, undergoes a “high-low-high” bimodal pattern regulating dynamic DNA methylation and gene expression during the naïve-formative-primed transitions. ZFP281 also safeguards DNA methylation in maintaining primed pluripotency. Our study demonstrates a previously unappreciated role for ZFP281 in coordinating DNMT3A/3B and TET1 functions to promote pluripotent state transitions.

**In Brief:** The naive, formative, and primed pluripotent states and their interconversions recapitulate pluripotency continuum during early development. Huang and colleagues investigated the transcriptional programs during successive pluripotent state transitions and revealed an essential role for ZFP281 in coordinating DNMT3A/3B and TET1 to establish the DNA methylation and gene expression programs during the transitions.

**Highlights:**

- ZFP281 activates *Dnmt3a/3b in vitro* in pluripotent stem cells and *in vivo* in epiblast.
- ZFP281 and TET1 undergo bimodal chromatin occupancy in pluripotent state transitions.
- Chromatin-binding of ZFP281 and TET1 depends on the formation of R-loops at promoters.
- ZFP281 is necessary for the establishment and maintenance of primed pluripotency.

## INTRODUCTION

Mammalian embryonic development involves DNA methylome remodeling with an initial drastic genome-wide DNA demethylation followed by the re-establishment of the methylation landscape within the pluripotent epiblast during the peri-implantation period.^1, 2^ Methylation of DNA cytosine to 5-methylcytosine (5mC) is an epigenetic modification associated with transcriptional repression.^3^ In mammalian cells, DNA methylation is established by the *de novo* DNA methyltransferases DNMT3A and DNMT3B and maintained by DNMT1 through cell division. Deletion of these enzymes in mice results in embryonic (*Dnmt1* or *Dnmt3b*) or postnatal (*Dnmt3a*) lethality.^4–6^ In contrast, DNA demethylation, mediated by the ten-eleven translocation (TET) dioxygenases, oxidizes 5mC to generate 5-hydroxymethylcytosine (5hmC) and its further oxidized species that can eventually be removed to become an unmethylated cytosine.^7^ The functional interaction between DNMT3 and TET shapes the mammalian DNA methylome and is essential for early development.^8–10^ DNMT3A and DNMT3B are recruited onto gene bodies through histone H3K36me2 and H3K36me3 marks along with gene transcription.^11, 12^ In contrast, the TET enzymes, which are primarily located at regulatory elements such as gene promoters and enhancers, form DNA hypomethylated valleys.^13–15^ However, how the TET enzymes are recruited to the tissue-specific gene regulatory regions remains elusive.^9^

DNA methylation and demethylation dynamics play critical roles in the regulation of gene expression during cell fate commitment and during early development coincident with the reorganization of the chromatin occupancy of pluripotency and tissue-specific transcription factors (TF) for target gene regulation. Pluripotency is a development continuum that encompasses a series of successive states,^16^ including the naive, formative and primed states of pluripotency that can be recapitulated by *in vitro* culture of embryonic stem cells (ESCs), epiblast-like cells (EpiLCs) and epiblast stem cells (EpiSCs), respectively.^17^ Pluripotency-associated TFs such as OCT4, SOX2, and NANOG are highly expressed in ESCs where they collaboratively inhibit lineage differentiation and preserve an undifferentiated state.^18^ Interestingly, while expression levels of OCT4 and SOX2 are relatively consistent in naive, formative and primed states, NANOG expression is reduced in the formative state, and its expression follows a biphasic pattern of ’on-off-on’ during the naive-formative-primed transition *in vitro* and *in vivo.*^19^ ZFP281 is a key partner of OCT4 in ESCs and during the naive-to-primed pluripotent state transition where it functions to reorganize enhancer landscapes^20–22^. We previously reported that ZFP281 coordinates opposing functions of TET1 and TET2 in epigenomic reconfiguration promoting the naive-to-primed pluripotent state transition *in vitro*^21^, corroborated by its essential and cell-autonomous roles in postimplantation epiblast development *in vivo*.^22, 23^

In this study, we further unravel the transcriptional and epigenetic mechanisms underlying the pluripotent state transitions by employing our *Zfp281* knockout (*Zfp281*KO) mouse model^22^ and a newly created degron system for rapid ZFP281 protein degradation in pluripotent stem cells (PSCs). We uncovered a previously unknown direct transcriptional control of *Dnmt3a* and *Dnmt3b* by ZFP281 in the early postimplantation at embryonic day 6.5 (E6.5) epiblast and in cultured PSCs recapitulating the naive-formative-primed transition and maintenance of primed pluripotency. Additionally, we discovered a bimodal pattern of “high-low-high” chromatin occupancy of ZFP281 and TET1, dependent on the formation of R-loops at ZFP281 targeted promoters, and an autoregulation of DNMT3A/3B during the pluripotent state transitions.

## RESULTS

### The transcriptional programs are dynamically regulated in PSCs and during early embryo development

The transition between successive pluripotent states within the epiblast *in vivo* can be recapitulated in *in vitro* cultured PSCs, including naive ESCs cultured in 2 inhibitors and LIF (2iL), metastable ESCs cultured in serum and LIF (SL), formative EpiLCs by adapting ESCs in Fgf2 and Activin culture (FA) for ∼2 days, and the primed EpiSCs in the long-term FA culture, as illustrated in Figure 1A. To understand the dynamic gene transcriptional programs associated with these distinct pluripotent states, we first compared the transcriptomes between metastable ESCs (SL) and formative EpiLCs (FA-D2) and identified 964 differentially expressed genes (DEGs, p- value<0.05, fold-change>2, Table S1). As expected for distinct formative and primed pluripotent states, most of them (62.3%, 601/964) were not among the DEGs by comparing ESCs and primed EpiSCs (Figure S1A-B). We then performed hierarchical clustering analysis for the 964 formative DEGs and identified four clusters (C1∼C4) with distinct expression patterns in ESCs, EpiLCs, and EpiSCs (Figure 1B). The C1 genes are mainly primed state-specific genes (e.g., *Zic2, Pitx2, Lefty1, Lin28a*) with progressively increased expression in EpiLCs and EpiSCs. In contrast, the C4 genes were mainly naive state-specific genes (e.g., *Tet2*, *Esrrb, Klf4/5, Prdm14*) with progressively decreased expression in EpiLCs and EpiSCs. We were particularly interested in the C2 and C3 cluster genes, which were upregulated (“formative-up” genes hereafter, e.g., *Dnmt3a/3b, Fgf5, Sall2, Otx2*) and down-regulated (“formative-down” genes hereafter, e.g., *Nanog, Etv4, Ccnd3, Pdgfa*) only in the formative state, respectively (Figure 1B). Gene ontology (GO) analysis of biological processes revealed that the C2 genes were involved in the “regulation of transcription, DNA-templated” and “DNA methylation”, while the C3 genes are involved in the “cell differentiation”, “nervous system development”, and “angiogenesis” (Figure S1C). Next, to understand the dynamic transcriptome change during the formative-to-primed transition, we further examined an intermediate state between formative and primed states, often referred to as the “late formative” state, by adapting ESCs in FA culture for 4 days (FA-D4) (Figure 1A). We plotted the average expression of C1∼C4 genes from different pluripotent states of PSCs (Figure 1C) and of corresponding embryonic stages^1^ (Figure 1D). Consistently, the formative-up (C2, e.g., *Dnmt3a/3b*) and formative-down (C3, e.g., *Nanog*) genes showed high and low expression levels, respectively, in E5.5-E6.5 epiblasts. Primed-specific genes (C1, e.g., *Zic2/5*) were highly expressed in E7.5 tissues, whereas naive-specific genes (C4, e.g., *Klf4*) were highly expressed in inner cell mass (ICM) of E3.5 and E4.0 embryos (Figure 1C-E).

**Figure 1.**
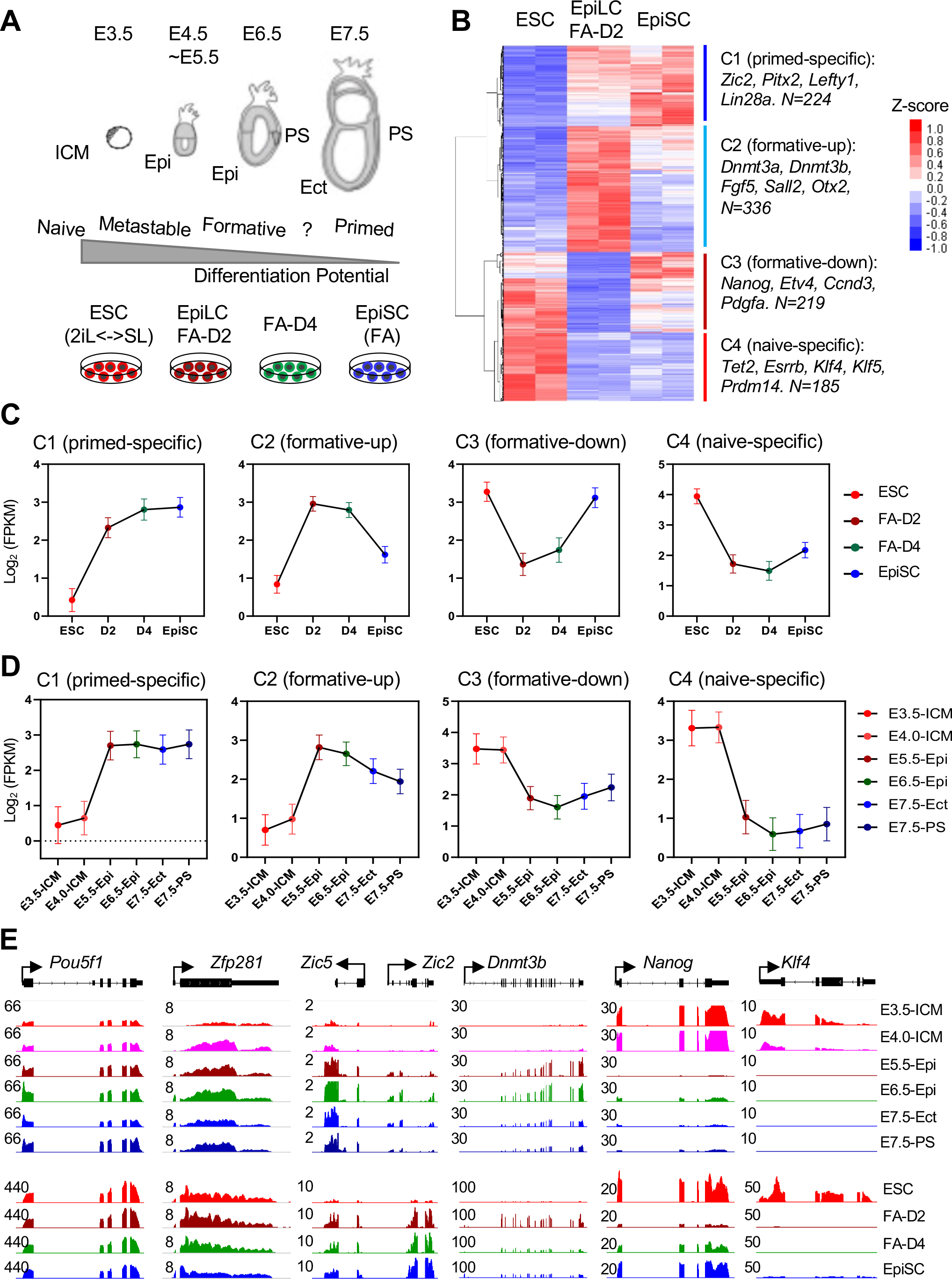
The dynamic gene expression during the pluripotent state transition. (A) An overview of mouse embryonic development from E3.5 to E7.5, representing the naive- formative-primed pluripotent state transition (top), recapitulated by the *In vitro* ESC-EpiLC-EpiSC transition (bottom). ESCs (in serum/LIF culture condition) were adapted in FGF2 and Activin (FA) culture for 2 days (FA-D2, EpiLCs) and 4 days (FA-D4), and further cultured under FA condition to generate stable primed EpiSCs. ICM, inner cell mass; Epi, epiblast; Ect, embryonic ectoderm; PS, primitive streak. “?” denotes a known intermediate state (i.e., FA-D4) between formative and primed pluripotency. (B) Heatmap depicting differentially expressed genes (DEGs) from RNA-seq analysis in ESCs, EpiLCs (FA-D2), and EpiSCs. Four clusters of genes with distinct expression patterns were identified. In each cluster, the total number and a few representative genes were labeled. (C-D) Average expression (by log2FKPM) of each gene cluster in different pluripotent stem cells (C) or embryo lineages (D). Line plots represent the mean expression value of each cluster with a 95% confidence interval. (E) RNA-seq tracks depicting expression of pluripotency-related genes in different pluripotent stem cells or embryo lineages (from our published study^1^). The numbers indicate the normalized RPM value of the tracks shown.

In contrast to dynamically expressed genes during pluripotent state transitions, other pluripotency factors, e.g., *Oct4 (Pou5f1) and Zfp281* (Figure 1E), were consistently expressed across all pluripotent states. We decided to focus on studying the molecular functions of ZFP281 in these pluripotent states and during the transition for the following reasons. First, we^21^ and others^24^ have shown that ZFP281 is essential for primed pluripotency, and *Zfp281* epiblast tissue specific mouse mutants manifested cell-autonomous developmental abnormalities at E6.0 and died E7.75,^22, 23^ spanning the formative-to-primed transition period *in vitro* (Figure 1A). However, our understanding of the mechanisms underlying this formative-to-primed transition is limited. Second, we have shown that ZFP281 interacts with TET1 and mediates transcriptional and posttranscriptional repression of TET2 in promoting primed pluripotency.^21^ We noted here that, like ZFP281, TET1 is also expressed across all pluripotent states, whereas TET2 is a “naive-specific” C4 gene that is downregulated in formative and primed states (Figure 1B). In contrast, TET3 is not expressed across any pluripotent states. Therefore, TET1 is the only functioning TET enzyme in formative and primed states.^25^ How ZFP281 might function with TET1 during the formative-to-primed transition has not been studied. Third, we also noted that *Dnmt3a/3b* are “formative-up” C2 genes, which are known to be activated by FGF signaling^26^ and upregulated in both formative and primed *in vitro* PSC cultures (Figures 1B and 1E, top) as well as in E5.5-E6.5 postimplantation embryos (Figure 1E, bottom). However, despite the well-recognized roles of DNA methylation in embryonic development,^1, 5, 27^ how the DNMT genes are transcriptionally activated in early embryos remains poorly understood. We thus decided to explore ZFP281 as a candidate factor regulating DNMT gene expression to fill this knowledge gap.

### *De novo* DNA methyltransferases are regulated by ZFP281 in postimplantation embryos

We first examined whether the DNMT genes are directly regulated by ZFP281 in the epiblast of developing embryos. From our RNA-seq analysis of wild-type (WT) and E6.5 *Zfp281*^-/-^ embryos,^22^ we found that mRNA expression levels of the DNMT3 family genes *Dnmt3a, 3b,* and *3l*, but not *Dnmt1*, are reduced in mutant embryos compared with WT embryos (Figure S2A). We also performed shRNA-mediated knockdown of *Zfp281* in ESCs and EpiSCs, and found that sh*Zfp281* reduced the expression of *Dnmt3a, 3b*, and *3l* genes, but not *Dnmt1* (Figure S2B), consistent with the *in vivo* results (Figure S2A). To assess the defects in *Zfp281* mutant embryos at the protein level, we performed single-cell quantitative immunofluorescence (qIF)^28^ and found that DNMT3A, 3B, and 3L proteins were significantly reduced in the *Zfp281*^-/-^ E6.5 epiblast compared with those in WT epiblast (Figure 2A-D). To examine whether DNA methylation is affected in the epiblast of *Zfp281* mutants, we applied STEM-seq^1^ (small-scale TELP-enabled methylome sequencing) for low-input genome-wide DNA methylome profiling in of WT, *Zfp281*^+/-^, and *Zfp281*^-/-^ E6.5 embryos. We found that DNA methylation at CG (mCG) sites decreased in the *Zfp281*^+/-^ embryos and further still in the *Zfp281*^-/-^ embryos relative to WT embryos at E6.5 (Figure 2E).

**Figure 2.**
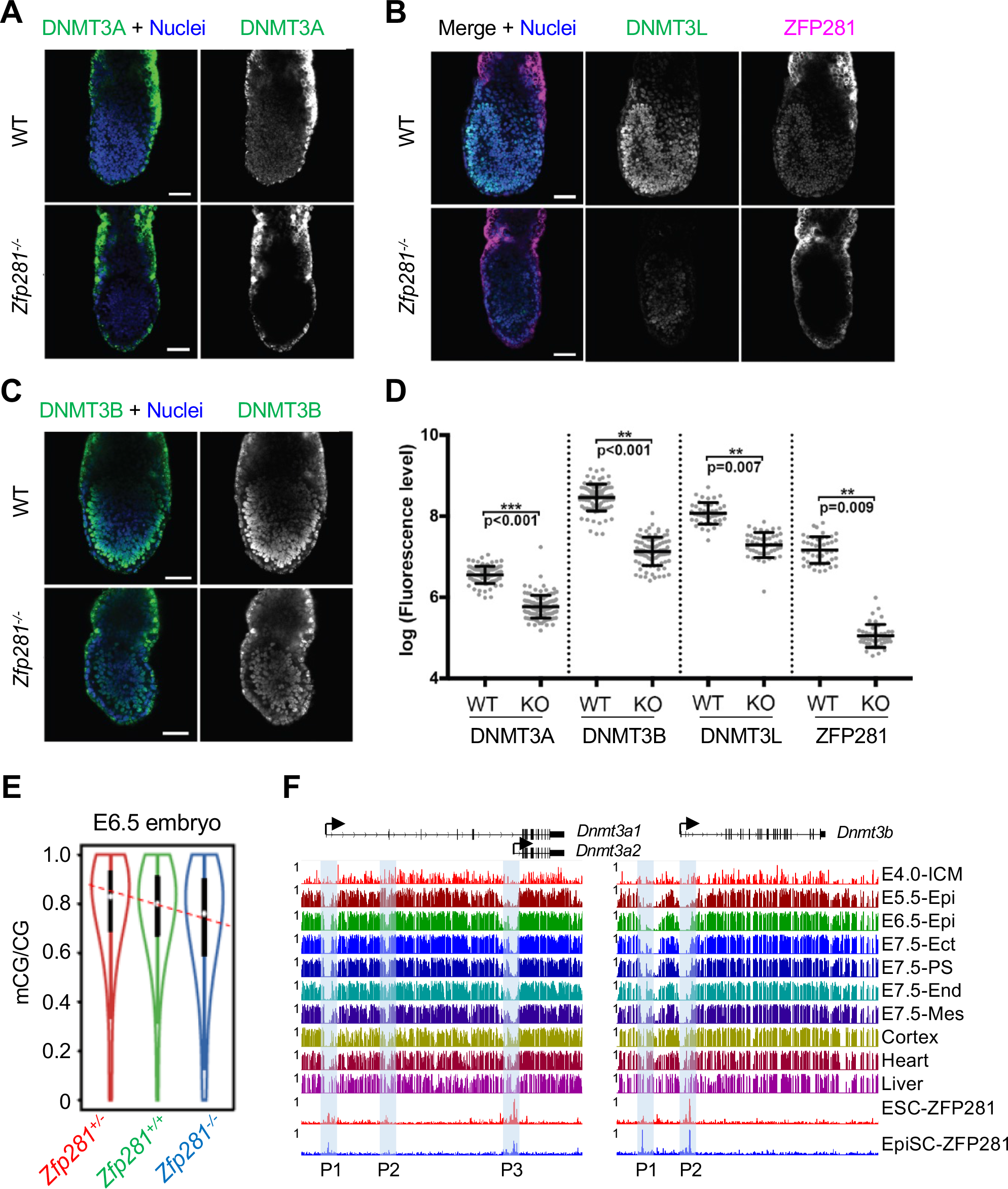
*De novo* DNA methyltransferases are regulated by ZFP281 in PSCs and postimplantation embryos. (A-C) Immunostaining of DNMT3A (A), DNMT3L and ZFP281 (B), and DNMT3B (C) in WT and *Zfp281*^-/-^ embryos (E6.5). (D) Fluorescent intensity of ZFP281 and DNMT3 family proteins was quantified using Imaris software. Each dot represents the mean corrected fluorescence level per epiblast cell. (E) The CG methylation (mCG) levels in the WT, *Zfp281*^+/-^, and *Zfp281*^-/-^ E6.5 embryos. (F) DNA-methylation landscapes in postimplantation embryonic lineages (E4.0-E7.5) and three adult tissues (from our published study^1^) and ZFP281 ChIP-seq tracks in ESCs and EpiSCs at *Dnmt3a* and *Dnmt3b* loci. The ZFP281 ChIP peaks (P1-P3) were labeled. The numbers indicate the mCG values and normalized RPM values of the tracks shown.

Chromatin immunoprecipitation (ChIP) followed by deep sequencing (ChIP-seq) of ESCs^21^ and EpiSCs^22^ and ChIP-qPCR analyses of ZFP281 in ESCs and EpiSCs revealed that ZFP281 directly binds to the regulatory elements at *Dnmt3a, 3b,* and *3l* loci (Figure S2C-D). Interestingly, the ESC and EpiSC ZFP281 binding peaks on *Dnmt3a* and *Dnmt3b* loci coincided with regions of DNA hypomethylation in postimplantation embryonic tissues (Figure 2F). Altogether, our results suggest that ZFP281 may transcriptionally activate *Dnmt3a/3b/3l* by direct binding at their regulatory loci (i.e., promoters) with DNA hypomethylation valleys during postimplantation development.

### ZFP281 directly activates DNMT3A/3B expression in controlling the transcription programs of formative and late formative pluripotent states

To further understand the transcriptional regulation of *Dnmt3a/3b* and address whether ZFP281 chromatin binding facilitates *Dnmt3a/3b* promoter demethylation during the peri-implantation embryonic development, we employed the *in vitro* pluripotent state transition model as a scalable alternative to the limited availability of embryonic tissues at these stages. Since loss of *Zfp281* affects the self-renewal of primed EpiSCs but not ESCs,^21^ we established a degron^29^ cell system for rapid, inducible and reversible ZFP281 protein degradation in ESCs (Figures 3A, two independent clones, #2 and #21, see details in Methods). Using *Zfp281*^degron^ ESCs, we confirmed that degradation TAG (dTAG) treatment could induce near-complete protein degradation within 2 hours (Figure 3A). Importantly, ZFP281 depletion by dTAG also decreased DNMT3A and DNMT3B expression in ESCs, while the removal of dTAG reintroduced their protein expression (Figure S3A). Of note, the *Dnmt3a* gene is transcribed from two alternative promoters (Figures 2F), resulting in two transcript isoforms: the long isoform *Dnmt3a1* and the short isoform *Dnmt3a2*, respectively.^13, 30^ The *Dnmt3b* gene also has multiple splice variants, but the *Dnmt3b* isoforms share the same promoter and have similar protein sizes. In ESCs, DNMT3A2 is predominantly expressed, while DNMT3A1 and DNMT3B isoforms are lowly expressed (Figure S3B). In the absence of dTAG, *Zfp281*^degron^ ESCs could be adapted in FA culture to derive EpiSCs, referred to as converted EpiSCs (cEpiSCs, Figure S3B). In contrast, *Zfp281*^degron^ ESCs could not be maintained in the FA culture for more than 3 passages in the presence of dTAG, consistent with our finding using *Zfp281*KO ESCs.^21^ We also examined ZFP281 function in the formative FA-D2 and late formative FA-D4 samples and found that depletion of ZFP281 by dTAG diminished the activation of DNMT3A and DNMT3B and delayed the reduction of naive-specific markers (e.g., KLF4, ESRRB) at both protein (Figures 3C and S3C) and RNA (Figure S3D) levels. More importantly, the global levels of DNA 5mC were decreased in FA-D2 (orange bars) and FA-D4 (light green bars) cells upon dTAG treatment measured by mass spectrometry quantification (Figure 3E) and DNA dot blot analysis of genomic DNA (Figure S3E). We also observed that DNA 5hmC was increased in the presence of dTAG (Figures 3E and S3E), likely due to the loss of DNMT3A/3B activity, leading to increased TET1 activity and DNA 5hmC.^13^

**Figure 3.**
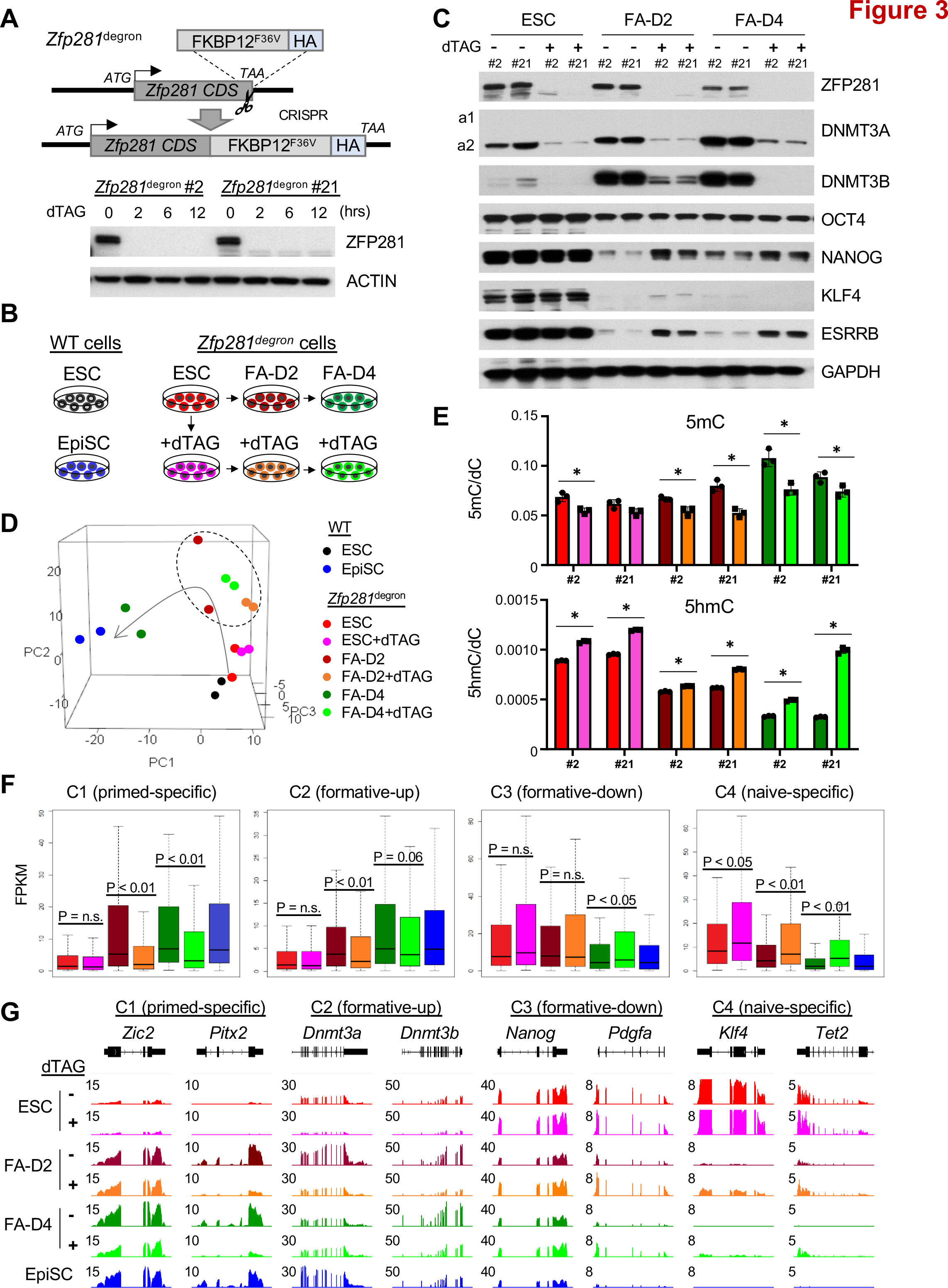
ZFP281 activates DNMT3A and DNMT3B in the pluripotent state transitions. (A) Schematic depiction of the *Zfp281*^degron^ knock-in (KI) strategy using CRISPR/Cas9 genome editing (the Scissor symbol). The HA-tagged FKBP12^F36V^ donor sequence was inserted right after the stop codon (TAA) of *Zfp281* CDS to create an in-frame fusion protein. Two independent clones (#2 and #21) of *Zfp281*^degron^ ESCs were derived. ZFP281 protein was depleted within 2 hours upon treatment of dTAG. (B) Schematic depiction of the *Zfp281*^degron^ ESCs in formative and primed transition. WT ESCs and EpiSCs were used as the reference states. *Zfp281*^degron^ ESCs were treated with dTAG for 2 days, and then both untreated and treated ESCs were adapted to FA culture condition without (top) or with (bottom) dTAG treatment. Samples were collected on Day 2 (FA-D2) and Day 4 (FA- D4). (C) Western blot analysis for *Zfp281*^degron^ ESCs, FA-D2, and FA-D4 cells. Two independent clones (#2, #21) were used with or without dTAG treatment. DNMT3A2 (a2) is the predominant isoform, and DNMT3A1 (a1) is faint in regular exposure. Another blot containing both DNMT3A isoforms by long exposure is shown in Figure S3C. (D) Principal component analysis (PCA) depicting RNA-seq samples of WT ESCs and EpiSCs, and *Zfp281*^degron^ cells in ESC-EpiLC-EpiSC transition with or without dTAG treatment. A transition trajectory was indicated by a curved arrow. (E) UHPLC-MS/MS quantification for 5mC (top) and 5hmC (bottom) intensities in the genomic DNA of *Zfp281*^degron^ PSCs with or without dTAG treatment. An intensity ratio of 5mC or 5hmC over deoxycytidine (dC) was measured. Experiments were performed in two independent clones (#2, #21) with technical triplicates; the p-value is from a two-tailed t-test, and ‘‘*’’ denotes p < 0.05. (F) Boxplots depicting expression of C1∼C4 genes in *Zfp281*^degron^ ESC-EpiLC-EpiSC transition with or without dTAG treatment. The P-value is from the Mann-Whitney test, and “n.s.” denotes statistically non-significant. (G) RNA-seq tracks depicting expression of representative C1∼C4 genes in *Zfp281*^degron^ ESC- EpiLC-EpiSC transition with or without dTAG treatment. The numbers indicate the normalized RPM values of the tracks shown. (D-G) Color codes match those in Panel B.

To understand how ZFP281 depletion and DNMT3A/3B downregulation affect gene transcription programs in distinct pluripotent states, we performed RNA-seq analysis of *Zfp281*^degron^ ESCs, FA-D2, and FA-D4 cells in the presence or absence of dTAG and compared them with embryo-derived WT ESCs and EpiSCs (Figure 3B). Principal component analysis (PCA) for the WT and *Zfp281*^degron^ ESCs, *Zfp281*^degron^ FA-D2 and FA-D4 cells without dTAG treatment, and WT EpiSCs revealed a trajectory of classic naive ESCs towards the formative/late formative and primed states (Figure 3D, the curved arrow). However, with dTAG, *Zfp281*^degron^ FA-D2 and FA-D4 samples deviated from the curve of transition. In addition, the dTAG-treated FA-D4 samples are far away from the untreated FA-D4 samples but relatively closer to the FA-D2 samples with or without dTAG treatment (Figure 3D, dashed circle), indicating a major defect in the formative-to-late formative transition upon ZFP281 depletion. We examined the expression levels of C1∼C4 genes (Figure 1B) in *Zfp281*^degron^ RNA-seq data and found that ZFP281 depletion compromised the activation of the C1 (primed-specific) genes (e.g., *Zic2, Pitx2*) and delayed the reduction of the C4 (naive-specific) genes (e.g., *Klf4, Tet2*) (Figure 3F-G). ZFP281 depletion also decreased the expression of the C2 (formative-up) genes in FA-D2 and FA-D4 samples (e.g., *Dnmt3a, Dnmt3b*) (Figure 3F-G). However, expression of the C3 (formative-down) genes was not affected in FA-D2 samples but increased in FA-D4 samples upon ZFP281 depletion (Figure 3F), suggesting that the C3 genes may be indirectly regulated by ZFP281, possibly by reduced DNMT3A/3B expression upon ZFP281 depletion.

The *de novo* DNA methylation established by DNMT3A/3B must be maintained by DNMT1 through cell division. Although DNMT1 is present within the epiblast at E6.5 and in PSCs, it is not regulated by ZFP281 (Figure S2A-B). To address whether the DNMTs are critical in the naive-to-primed pluripotent state transition, we employed WT, *Dnmt1*-KO, *Dnmt3a/3b*-DKO, and *Dnmt1/3a/3b*-TKO ESCs and performed ESC-to-EpiSC differentiation. We were able to convert *Dnmt1*-KO and *Dnmt3a/3b*-DKO ESCs to cEpiSCs, which is consistent with the finding that *Dnmt3a/3b* single or double KO mice appeared to develop normally until E8.5.^5, 27^ However, we failed to convert *Dnmt1/3a/3b*-TKO ESCs to cEpiSCs (Figure S3F-G), consistent with the G2/M cell cycle arrest and enhanced cell death during epiblast differentiation of these TKO cells.^31^ The apparent discrepancy in the establishment of the primed pluripotent state between *Zfp281*KO^21^ and *Dnmt3a/3b*-DKO ESCs could be due to the engagement of additional key factors downstream of ZFP281 in this state transition^24^ (see Discussion).

### Dynamic chromatin occupancy of ZFP281/TET1 and feedback control of DNMT3A/3B in the pluripotent state transitions

Given that activation of DNMT3 family genes depends on the transcriptional activity of ZFP281 (Figures 2 and 3) and that ZFP281 can recruit TET1 at ZFP281-bound promoters for targeted DNA demethylation at certain loci,^21^ we asked how ZFP281 might coordinate both classes of DNA epigenetic regulators to regulate downstream target genes during the pluripotent state transitions. We performed ChIP-seq analysis of ZFP281 by antibody pulldown in WT ESCs, FA-D2, FA-D4 cells, and EpiSCs and by HA-tag pulldown in *Zfp281*^degron^ ESCs, FA-D2, FA-D4 cells (Figures 3A and 4A). The ChIP-seq data obtained from the two pulldowns showed good correlation at different pluripotent states (Figure S4A), demonstrating the quality of the dataset. First, we compared the ZFP281 peaks identified in WT ESCs and EpiSCs and observed a genome-wide rearrangement of chromatin-bound ZFP281 (Figure 4B). Motif analysis for the ESC-only, EpiSC-only, and ESC/EpiSC-shared peaks of ZFP281 identified a consensus G-rich motif (Figures 4C and S4B), suggesting ZFP281 actively binds to its target loci in respective pluripotent states. When plotting the ZFP281 ChIP intensity at all its peak regions (N=12,732), we were surprised to note that the average intensity in the FA-D2 samples was significantly lower than that of the other states (Figure 4D). Next, we performed TET1 ChIP-seq in the same cell states (Figure 4A). Interestingly, we observed low TET1 ChIP intensity at the ZFP281 peaks in the FA-D2 samples (Figure 4E). However, such a trend was not observed by plotting TET1 ChIP intensity at all transcription start sites (TSSs) (Figure S4C), reinforcing a ZFP281-dependent recruitment of TET1 at the ZFP281 target sites during the naive-to-formative transition. When comparing the ZFP281 peak intensities in different PSC states, we found that the intensities of most peaks were decreased (N=3,804, false discovery rate < 0.05) in the ESC-to-FA-D2 transition. In contrast, the intensities of most peaks were increased (N=1,695) in the FA-D2-to-FA-D4 transition, and the intensities of fewer peaks were changed when comparing the ESC and FA-D4 states (Figure 4F, H), highlighting a major regulatory event happening during the formative to late formative state transition. We then compared the 3,084 and 1,695 significantly altered ZFP281 peaks and found that many of the same peaks (N=945, Chi-square test, *P* = 2.2 e^-16^) decreased in ESC-to-FA-D2 transition and then recovered in FA-D2-to-FA-D4 transition (Figure 4G). GO analysis for the target genes of 945 ZFP281 peaks (TSS < 5K) revealed that they are involved in the “regulation of cell cycle”, “multicellular organism development”, “chromatin organization”, and “cell differentiation” (Figure S4D). Together, our data suggest that chromatin-bound ZFP281 and TET1 at a subgroup of ZFP281 peaks exhibited a bimodal “high-low-high” occupancy pattern during the naive-formative- late formative transition, which was maintained high thereafter till the primed EpiSC state (Figure 4I). The rebounced chromatin occupancy of ZFP281 and TET1 in the late formative state is likely a key epigenetic event via DNA hypomethylation to reactivate those ZFP281 target genes that are downregulated in the formative state (Figure 4H, the C3 genes).

**Figure 4.**
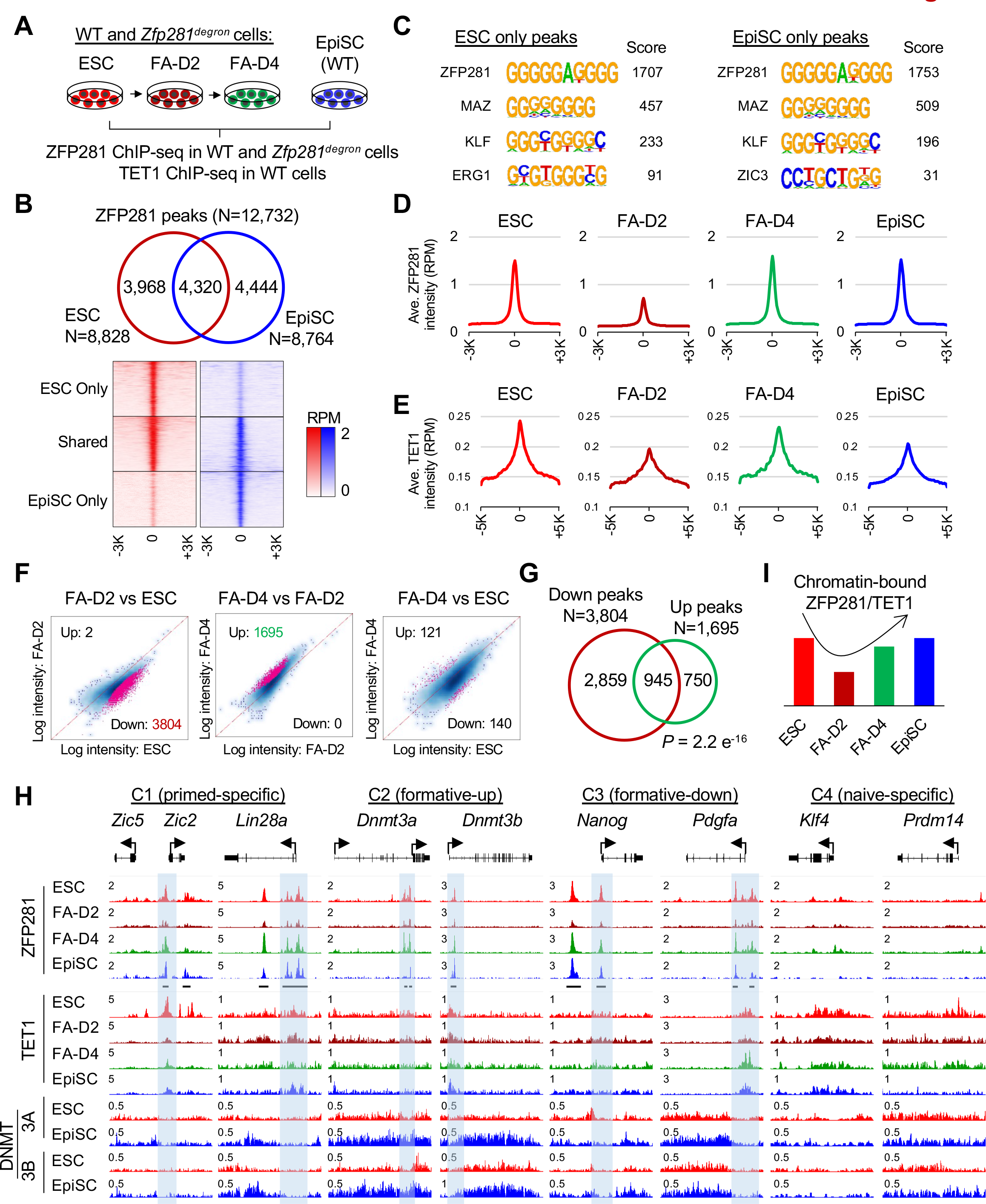
Dynamic chromatin occupancy of ZFP281/TET1 and feedback transcriptional control of DNMT3A/3B in the pluripotent state transitions. (A) Schematic depiction of the ZFP281 and TET1 ChIP-seq analysis using WT and *Zfp281*^degron^ PSCs. ZFP281 ChIP-seq was performed in WT and *Zfp281^degron^* ESCs, FA-D2, FA-D4 cells, and WT EpiSCs. TET1 ChIP-seq was performed in WT ESCs, FA-D2, FA-D4 cells, and EpiSCs. (B) Overlap of the identified ZFP281 peaks (top) and intensity heatmap (bottom) from ZFP281 ChIP-seq analysis in WT ESCs and EpiSCs. (C) The top enriched motifs identified from the ESC-only and EpiSC-only ZFP281 peaks. (D) Mean intensity plots depicting ZFP281 ChIP-seq intensity at ZFP281 peak regions (within ± 3K bp around the peak center). ZFP281 ChIP-seq data in WT and *Zfp281^degron^* ESCs, FA-D2 and FA-D4 cells were merged. (E) Mean intensity plots depicting TET1 ChIP-seq intensity at ZFP281 peak regions (within ± 5K bp around the peak center) in WT ESCs, FA-D2 and FA-D4 cells, and EpiSCs. (F) Scatterplots depicting the distribution of ZFP281 peak intensities by comparing FA-D2 vs. ESC (left), FA-D4 vs. FA-D2 (middle), and FA-D4 vs. ESC (right) ChIP-seq data. Peaks with significantly different intensities were determined by FDR < 0.05. (G) Overlap of the significantly decreased ZFP281 peaks (N=3,804) by comparing FA-D2 vs. ESC data and significantly increased peaks (N=1,695) by comparing FA-D4 vs. FA-D2 data. The p-value is from the Fisher’s extract test. (H) ChIP-seq tracks depicting the intensities of ZFP281, TET1, DNMT3A, and DNMT3B at representative C1∼C4 genes in different PSCs. The identified ZFP281 peaks were labeled on the bottom of the ZFP281 tracks. The numbers indicate the normalized RPM value of the tracks shown. (I) Schematic depiction of the chromatin-bound ZFP281 and TET1 in ESCs, FA-D2, FA-D4 cells, and EpiSCs.

Next, we asked whether other pluripotency-associated TFs or epigenetic regulators are dynamically distributed along with ZFP281 during pluripotent state transitions. Previously we showed that ZFP281 interacts with the pluripotency TF OCT4 and histone acetyltransferases P300/P400 in ESCs and EpiSCs.^21, 22^ From a published dataset of ESC-to-FA-D2 transition,^20^ we found that ChIP intensities of OCT4 and P300 were decreased along with ZFP281 (at 3,804 ZFP281 peaks with decreased intensities) (Figure S4E). From another published dataset of a 3- day EpiSC-to-ESC reprogramming experiment,^32^ we observed a similar pattern, such that OCT4 and NANOG showed decreased ChIP intensities from day 0 to day 1, and then increased intensities at days 2 and 3 at the ZFP281 peaks, as well as at all the identified OCT4 or NANOG peaks (Figure S4F-G). Together, these data support our finding that ZFP281, as a component of a transcriptional complex containing OCT4 and P300 with consistent expression levels in different pluripotent states (Figures 1E and 3C), follows a bimodal pattern of chromatin occupancy with “high-low-high” binding intensities during the naïve-formative-primed transitions.

Since DNMT3A and DNMT3B are highly activated in the primed state (Figure S3B), we also performed DNMT3A/3B ChIP-seq in WT EpiSCs and compared the data with the ChIP-seq data in ESCs.^33^ We found that DNMT3A/3B mainly bind across gene bodies to deposit DNA 5mC modification in ESCs and EpiSCs (Figure S4H, “All genes”), while hypomethylation status at promoters was protected by TET1 and additional factors (i.e., ZFP281) in ESCs and EpiSCs as well as during the pluripotent state transitions (Figure 4H, gene promoters with ZFP281 peaks in C1∼C3). Interestingly, when examining the DNMT3A and DNMT3B ChIP intensities across gene bodies of C1∼C4 cluster genes associated with the state transitions (Figure 1B), we found that only C1/C2 but not C3/C4 genes displayed higher DNMT3A/3B-binding intensities in EpiSCs than ESCs (Figure S4H), likely because C1/C2 genes are transcriptionally activated during the naive-to-primed transition (Figure 1B) and that DNMT3A/3B bind to gene bodies that undergo active transcription.^12, 33^ Since *Dnmt3a* and *Dnmt3b* are among the C2 genes, our data also suggest a transcriptional autoregulation of Dnmt3a and Dnmt3b (Figure 4H), downstream of their activation by ZFP281, during pluripotent state transitions.

### ZFP281 chromatin association depends on the formation of R-loops

We further explored the mechanism underlying the dynamic ZFP281 chromatin association during the pluripotent state transitions. We showed that ZFP281 mainly binds to DNA with a high G-rich motif (Figures 4C and S4B). It is well-established that active transcription on the G-rich (GC skew) DNA template strands is prone to R-loop formation.^34^ Interestingly, R-loops are also associated with promoters occupied by TET1 bearing 5hmC modification in ESCs^35^ and protect DNA from de novo methylation.^34^ Therefore, we investigated the relationship between ZFP281/TET1 chromatin binding and the formation of R-loops at ZFP281 targeted promoters.

We processed publicly available R-loop profiling datasets in ESCs using MapR,^36, 37^ a modified CUT&RUN-based method that uses a catalytically inactive RNase H1 (gene name: *Rnaseh1,* an endogenous gene that specifically recognizes and digests R-loop) fused to micrococcal nuclease (MNase). Overall, R-loop intensities at all gene promoters are associated with the activities of transcription (Figure 5A). We compared the R-loop enrichment at promoters of genes with low (FPKM<1), intermediate (1<FPKM<10), and high (FPKM>10) expression in ESCs and found that ZFP281 target genes have higher R-loop intensities at promoters than the non-target genes in each group (Figure 5B). Next, we plotted the R-loop intensities at the promoters of the C1∼C4 cluster genes. Although expression of the C1/C2 (i.e., primed-specific/formative-up) genes is extremely low in ESCs (Figure 1B-C), there are considerable R-loops enriched at their promoters (Figure S5A). The C3/C4 (i.e., formative-down/naive-specific) genes are highly expressed in ESCs (Figure 1B-C), and consistently, R-loops are also enriched at their promoters (Figure S5A). Therefore, our data indicate that the R-loop formation is relatively pervasive at promoters of genes regardless of their expression levels.

**Figure 5.**
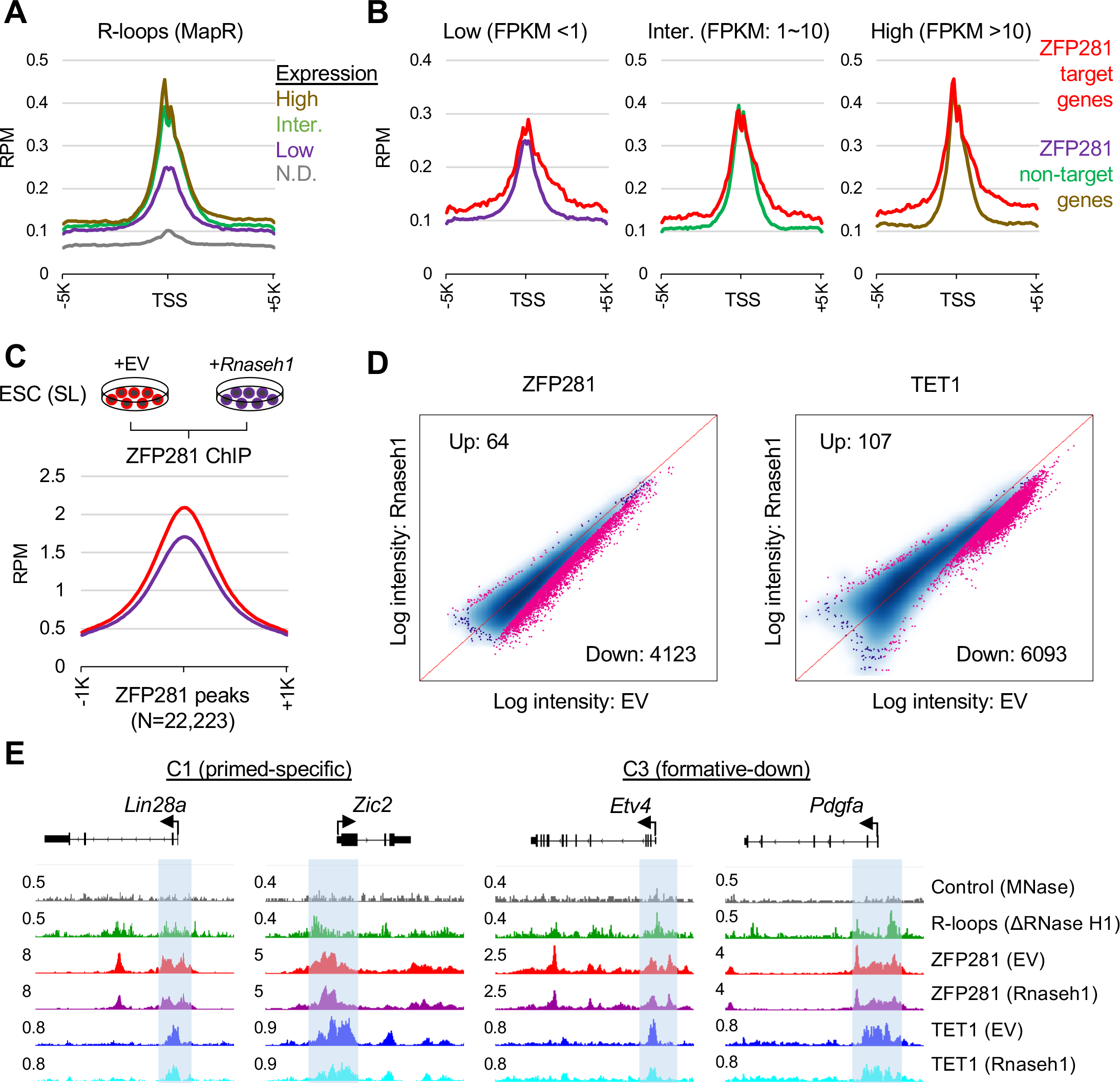
ZFP281 chromatin association depends on the formation of R-loops. (A) R-loop intensity by MapR (from a published study^37^) enriched at gene transcription state sites (TSS ± 5K bp). Genes were grouped by their expression values in ESCs by the fragments per kilobase per million reads (FPKM) values: Low (FPKM <1); Intermediate (FPKM: 1∼10); High (FPKM >10); N.D., not detected. (B) R-loop intensity enriched at TSSs (± 5K bp) of ZFP281 ChIP target genes (red curve) versus the rest of the non-target genes in each expression group, as shown in Panel A. (C) Mean intensity plots depicting ZFP281 ChIP intensity enriched at ZFP281 peaks (within ± 3K bp around the peak center) in ESCs with empty vector (EV) treatment or *Rnaseh1* overexpression. (D) Scatter plots depicting the distribution of ZFP281 (left) and TET1 (right, from a published study^38^) ChIP intensities at ZFP281 peaks (N=22,333) in ESCs with EV treatment or *Rnaseh1* overexpression. Peaks with significantly different intensities were determined by FDR < 0.05. (E) R-loop intensity by MapR, and ZFP281, TET1 ChIP-seq tracks depicting the intensities at the ZFP281/TET1 target promoters (labeled with blue shadow). The numbers indicate the normalized RPM value of the tracks shown.

Finally, we asked whether ZFP281 binding at chromatin depends on the formation of R-loops. To this end, we performed ZFP281 ChIP-seq analysis in ESCs with either empty vector (EV) or *Rnaseh1* overexpression (Figure 5C). Since TET1 can be recruited to chromatin by ZFP281, we also processed published TET1 ChIP-seq data with the same treatment of *Rnaseh1* overexpression in ESCs.^38^ Interestingly, both ZFP281 and TET1 ChIP intensities decreased at the ZFP281 peak regions upon *Rnaseh1* overexpression (Figures 5C-D and S5B), suggesting that chromatin association of ZFP281 and TET1 depends on the formation of R-loops. GO analysis for the R-loop-sensitive ZFP281 target genes revealed that they are involved in the “regulation of transcription, DNA-templated”, “chromatin organization”, “cell differentiation”, “nervous system development”, and “WNT signaling pathways” (Figure S5C). These R-loop-sensitive ZFP281 target genes encompass all the 4 cluster genes and examples for a few C1 (primed-specific) and C3 (formative-down) genes are given (Figure 5E). These selected C1 and C3 genes are upregulated during the formative-to-primed pluripotent state transition and are direct target genes of ZFP281/TET1 with coincident R-loop formation and binding of ZFP281 and TET1 at targeted promoters (shaded region in Figure 5E). Together, our data suggest a potential R-loop-dependent positive feedback regulation between the transcriptional activity at GC-skew, DNA hypomethylated promoters and ZFP281/TET1 chromatin binding in pluripotent state transitions.

### ZFP281 safeguards the homeostasis of DNA methylation and demethylation in maintaining primed pluripotency

Having established the critical roles of ZFP281 in coordinating DNMT and TET functions during the naive-formative-primed pluripotent state transitions, we finally decided to investigate the mechanism by which ZFP281 maintains primed pluripotency, considering the previously observed incompatibility between ZFP281 loss and EpiSC self-renewal by ourselves^21^ and others^24^. We took advantage of the *Zfp281*^degron^ cEpiSCs (Figure S3B) and treated them with dTAG for 2 days (D2) and 4 days (D4), followed by RNA-seq analysis (Figure 6A). *Zfp281*^degron^ cEpiSCs could not be maintained for 3 passages with persistent dTAG treatment (data not shown). There were more DEGs identified at D4 than D2 of dTAG treatment (comparing D4 vs. D0 and D2 vs. D0) but with a consistent trend of transcriptome change at the two time points (Figure S6A-B), suggesting that transcriptional changes upon ZFP281 depletion are enhanced with a more prolonged dTAG treatment. We performed hierarchical clustering analysis and identified the up- and down- regulated DEGs upon ZFP281 depletion in cEpiSCs (Figure 6A, Table S2). Genes associated with lineage development (e.g., *Lefty1, Lefty2, Lin28a*), de novo DNA methylation (i.e., *Dnmt3a/3b*), WNT signaling pathway and downstream targets (e.g., *Wnt3, Wnt8a, Lrp5, Nkd1, Nkd2, Ccnd1, Ccnd2*), and somatic differentiation (e.g., HOX cluster genes) were downregulated upon ZFP281 depletion (Figures 6A-C). In contrast, most naive pluripotency genes (e.g., *Esrrb, Zfp42, Tet2, Klf2, Klf4, Prdm14*), DNA demethylation enzymes *Tet1/Tet2*, and factors that are important for primed pluripotency (e.g., *Fgf5, Otx2, Zic3*), were upregulated upon ZFP281 depletion (Figures 6A-C). These results indicate that ZFP281 depletion in cEpiSCs disrupts the primed pluripotency gene expression programs.

**Figure 6.**
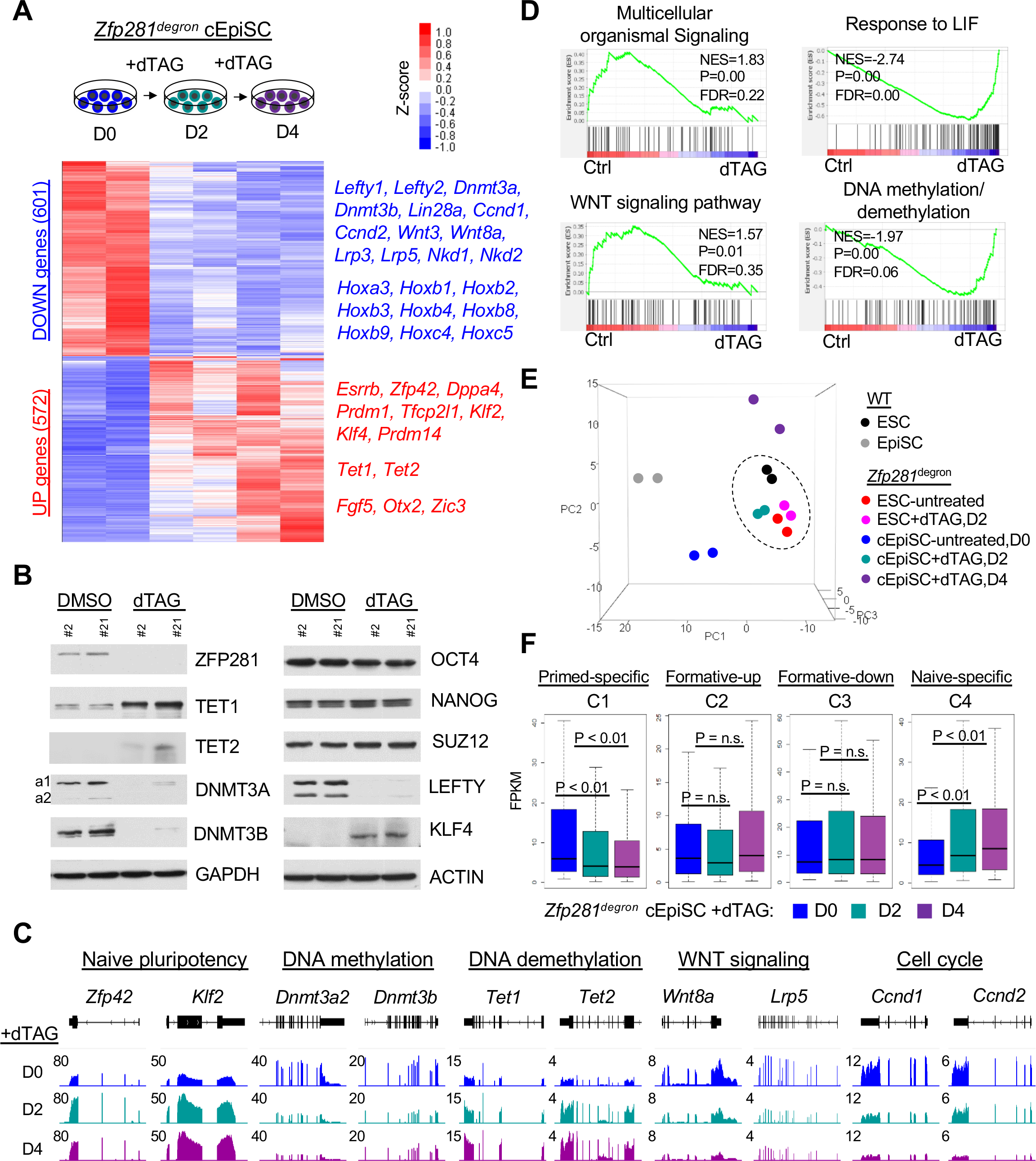
ZFP281 safeguards the homeostasis of DNA methylation and demethylation in maintaining primed pluripotency. (A) Heatmap depicting DEGs from RNA-seq analysis in *Zfp281*^degron^ cEpiSCs before (D0) and after dTAG treatment for 2 (D2) and 4 (D4) days. Representative up- and down-regulated genes were shown on the right side of the heatmap. (B) Western blot analysis for *Zfp281*^degron^ cEpiSCs with control DMSO or dTAG treatment (4 days). (C) RNA-seq tracks depicting expression of genes in *Zfp281*^degron^ cEpiSCs with dTAG treatments. The representative genes in different functional groups were labeled. The numbers indicate the normalized RPM value of the tracks shown. (D) Gene set enrichment analysis (GSEA) for the enriched oncology gene sets by comparing the control WT EpiSCs and untreated *Zfp281*^degron^ cEpiSCs (Ctrl) versus the dTAG-treated (D2, D4) *Zfp281*^degron^ cEpiSCs. Normalized enrichment score (NES), P-value, and false discovery rate (FDR) for the enrichment were indicated with the enrichment plot. (E) Principal component analysis (PCA) depicting RNA-seq samples of WT ESCs and EpiSCs, and *Zfp281*^degron^ cEpiSCs with dTAG treatments. A dash line circle contains WT and *Zfp281*^degron^ ESCs, as we all *Zfp281*^degron^ cEpiSCs with 2 days of dTAG treatment. (F) Boxplots depicting expression of C1∼C4 genes in *Zfp281*^degron^ cEpiSCs with dTAG treatments. The p-value is from the Mann-Whitney test, and “n.s.” denotes statistically non-significant.

To gain a global view of the transcriptional changes, we performed gene set enrichment analysis (GSEA) and GO analysis for the RNA-seq data. These analyses revealed that WNT signaling pathway and DNA methylation/demethylation are among the most significantly perturbed upon ZFP281 depletion (Figures 6D and S6C). Consistently, 5mC and 5hmC levels, measured by DNA dot blot (Figure S6D) and quantified by mass spectrometry (Figure S6E), were decreased and increased, respectively, upon ZFP281 depletion, likely due to decreased DNMT3A/3B and increased TET1/2 in cEpiSCs (Figure 6A-C). We also performed PCA analysis for the RNA-seq data of WT ESCs and EpiSCs, and *Zfp281*^degron^ ESC and cEpiSCs with dTAG treatments, revealing that the transcriptome of D2+dTAG *Zfp281*^degron^ cEpiSCs was closer to that of WT and *Zfp281*^degron^ ESCs with or without dTAG (Figure 6E, dashed circle), indicating a short-term ZFP281 depletion in cEpiSCs was driving the reversion of primed transcriptome to an ESC state. However, extended dTAG treatment caused the transcriptome of D4+dTAG *Zfp281*^degron^ cEpiSCs (Figure 6E, purple dots) to deviate away from the WT ESCs (black dots), *Zfp281*^degron^ ESCs with or without dTAG (pink and red dots), WT EpiSCs (grey dots), or untreated *Zfp281*^degron^ cEpiSCs (blue dots), suggesting that long-term Zfp281 depletion in cEpiSCs would cause the loss of primed pluripotency without reverting to an ESC state. Furthermore, when examining the expression of C1∼C4 cluster genes (Figure 1B) in the *Zfp281*^degron^ cEpiSCs with dTAG treatments (D0, D2, D4), we found that ZFP281 depletion decreased the expression of C1 (primed-specific) genes and increased the C4 (naive-specific) genes (Figure 6F). However, expression of C2 (formative-up) and C3 (formative-down) genes was not affected by ZFP281 depletion, suggesting that ZFP281’s transcriptional regulation of C2 and C3 genes is critical for the ESC-EpiLC-EpiSC transition but dispensable for EpiSC self-renewal/maintenance (Figures 3F and 6F). Together, these results demonstrate that, while distinct transcription programs are regulated by ZFP281 for the maintenance versus the establishment of the primed state of pluripotency, the regulation of DNA methylation/demethylation homeostasis is likely a conserved function of ZFP281 essential for both these processes.

## DISCUSSION

Pluripotency is highly dynamic and is under a tight transcriptional and epigenetic control with gene activation and repression generally corresponding to DNA demethylation and methylation in the context of different states of pluripotency. Studies of the gene expression programs of naive and metastable ESCs, formative EpiLCs, primed EpiSCs, and their interconversions have enriched our molecular understanding of pluripotency progression and cellular reprogramming.^17, 20, 32, 39, 40^ In this study, we established a central role of ZFP281 in dynamically regulating gene expression via the coordination of DNMT3 and TET1 functions in pluripotent state transitions. We identified 4 groups of genes whose expression is differentially regulated during the naive-to-formative-to-primed cell state transitions (Figure 7A) by a distinct bimodal pattern of ZFP281 and TET1 chromatin occupancy with an increase of DNA methylation (Figure 7B). Moreover, we identified a FA-D4 “late formative” state of pluripotency where the ZFP281/TET1 chromatin binding patterns are like the primed state (Figures 4D-E and 7B), although the gene expression profiles are closer to the FA-D2 formative state (Figures 1C and 7A). The relatively narrow bimodal ZFP281/TET1 chromatin occupancy (Figure 7B) and broader biphasic C2/C3 gene expression (Figure 7A) patterns support that chromatin reorganization by TFs and epigenetic regulators is an earlier event than the gene expression changes during the pluripotent state transitions. Detailed molecular characterization and functional studies using *Zfp281*KO mouse embryos and *Zfp281*^degron^ PSCs establish a complex interplay among ZFP281, TET1, and DNMT3A/3B (Figure 7C). First, ZFP281 and TET1 form a partnership to co-occupy at target genes (Figure 4D-E) and transcriptionally activate them including *Dnmt3a/3b* genes (Figure 2A-D, S2A-B, and Figure 3C, S3A,D). Second, DNMT3A/3B transcriptionally autoregulate their own genes (Figure 4H and S4H). Third, DNMT3A/3B and TET1 are functionally antagonizing each other in 5mC/5hmC modifications (Figure 3E, S3E and Figure S6D-E), likely due to their physical competition in chromatin occupancy (Figure 7C), as reported in ESCs.^13^ Our study deepens our understanding of the transcriptional and epigenetic mechanisms underlying pluripotent state transitions and early development.

**Figure 7.**
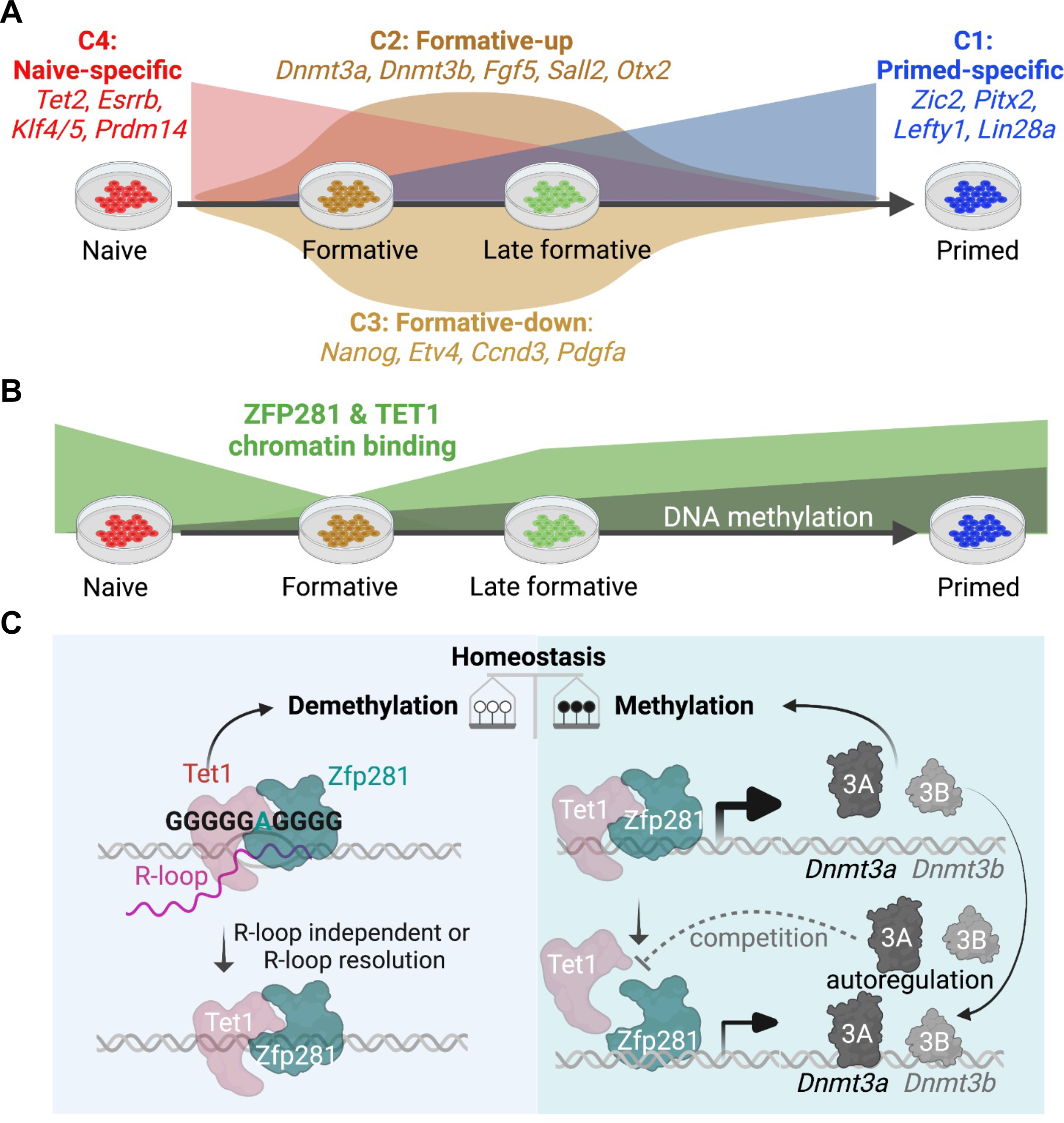
A summary model of ZFP281 functions in controlling transcription and DNA methylation landscapes in pluripotent state transitions. (A) Depiction of the 4 clusters of genes (C1/primed-specific, C2/formative-up, C3/formative-down, and C4/naive-specific) with distinct expression patterns during the naive, formative, late formative, and primed pluripotent state transitions. The representative genes are shown. (B) Depiction of the bimodal “high-low-high” pattern of ZFP281 and TET1 chromatin occupancy and an increase of DNA methylation during the naive, formative, late formative, and primed transitions. Depiction of the interplay among ZFP281, TET1, and DNMT3A/3B for the DNA methylation/demethylation homeostasis during the pluripotent state transitions. Left: ZFP281- TET1 chromatin co-occupancy at ZFP281 target sites is dependent on R-loops. The presumed function of ZFP281 in R-loop resolution and potential R-loop independent chromatin binding by ZFP281 and/or TET1 are also shown. Right: ZFP281 and TET1 transcriptionally activate *Dnmt3a* and *Dnmt3b* in E6.5 epiblast and different PSCs (ESCs, EpiLCs, EpiSCs), whose translation products (DNMT3A and DNMT3B) bind to their own gene bodies for autoregulation. The potential competition between TET1 and DNMT3A/3B, reported previously in ESCs^13^ and indicated with dashed inhibition lines, would lead to downregulation of ZFP281-regulated genes including *Dnmt3a/3b* (and thus reduced 5mC levels), which in turn relieves TET1 from the competition, leading to heightened TET1 chromatin binding (and thus increased 5hmC levels).

Naive ESCs maintained under 2i and LIF culture exhibit genome-wide hypomethylation and an open chromatin landscape. However, hypomethylation in naive ESCs also leads to eroded genomic imprints and chromosomal abnormalities.^41, 42^ In contrast, metastable SL ESCs are constrained from differentiation by serum and LIF with considerable expression levels of *Dnmt3a/3b/3l* and *Tet1/2* genes to preserve genomic stability^26^, and are thus functionally naive. Using SL ESCs as a starting point for pluripotent state transitions, we demonstrate that ZFP281 directly activates DNMT3A and DNMT3B in different pluripotent states. DNMT3L, a cofactor of DNMT3A/3B, is also activated by ZFP281 in mouse epiblast (Figure 2D), but such regulation could not be validated *in vitro* due to the low level of expression of *Dnmt3l* in formative and primed states. In the epiblast of E6.5 mouse embryos, which corresponds to the primed state of pluripotency, *Dnmt1* gene expression is unaffected by *Zfp281* KO. However, DNMT3A/3B/3L gene and protein expressions are compromised, although not abrogated, by *Zfp281* KO (Figures 2A-D and S2A), thereby explaining the mild decrease of DNA methylation in E6.5 *Zfp281*KO embryos (Figure 2E). However, the fact that we could establish primed cEpiSCs from *Dnmt3a/3b-* DKO ESCs, but not *Zfp281*KO^21^ ESCs, suggests that additional transcriptional or epigenetic regulators downstream of ZFP281 are necessary for the establishment of primed pluripotency. Indeed, the histone H3 lysine 9 (H3K9) methyltransferase EHMT1 and zinc finger TF ZIC2 were reported two critical functional players acting downstream of ZFP281 both in driving exit from the ESC (naive) state and in restricting reprogramming of EpiSCs (primed) to an ESC state^24^. In addition, we have shown that ZFP281 activates *miR-302/367* to repress TET2 expression for establishing and maintaining primed pluripotency.^21^ Of note, the defects observed in *Zfp281*KO embryos include failure of distal visceral endoderm/anterior visceral endoderm (DVE/AVE) specification, migration, and anterior-posterior (A-P) axis formation at E6.5,^22^ which may or may not be directly linked to the DNA epigenetic regulation by ZFP281 revealed in this study. However, the roles of ZFP281 in regulating DNMT3 and TET1 activities at gastrulation (E6.5-E8.5) are possible, based on a single-cell RNA-seq dataset of mouse gastrulation (E6.5-E8.5)^43^ which reveals that *Zfp281,* but not *Oct4* or *Nanog,* is broadly expressed together with *Dnmt3a/3b* and *Tet1* in the epiblast and in different lineage-specific precursors from E6.5 onwards (Figure S7A- B). Therefore, we conclude that ZFP281 is an important TF orchestrating the transcription and DNA methylation landscapes during pluripotent state transitions and early embryonic development.

R-loops have long been considered an accidental by-product of transcription, although their roles in opening double-stranded DNA structures to maintain active transcription are also recognized.^44^ Depletion of R-loops by overexpressing *Rnaseh1* in ESCs mildly impedes ESC differentiation.^45^ We established R-loop dependent chromatin occupancy of ZFP281 and TET1 in ESCs (Figure 5D-E), which may partly explain the function of R-loops in ESC differentiation. R-loops are reported to be enriched at TET1-bound promoters enabling elevated transcription, although such TET1/5hmC/R-loop-rich loci are prone to DNA damage.^35^ Interestingly, a recent study also suggests that ZFP281 is a critical factor for R-loop resolution through recruitment of BRCA2,^46^ a DNA damage repair associated factor that can further recruit RNA helicase DDX5 to promote R-loop resolution.^47^ Therefore, ZFP281 may have dual functions related to R-loops: On one hand, the formation of R-loops maintains ZFP281 and TET1 chromatin binding; on the other hand, ZFP281 employs the R-loop resolution mechanism to protect genome stability. Future studies are warranted to understand the functional interplay between R-loops and ZFP281 for genome stability during these critical cell state transitions.

## Limitations of the study

Our current study demonstrates that both DNMT3A and DNMT3B are transcriptionally activated by ZFP281 during the pluripotent state transition. However, the two proteins are known to have unique functions in ESCs and early development.^27, 33, 48, 49^ Future studies are needed to distinguish the functions and targets of DNMT3A versus DNMT3B (and possibly the isoforms associated with each gene) in different PSCs and during pluripotent state transitions. In addition, DNMT1 was reported to possess *de novo* DNA methylation activity in mouse oocytes,^50^ but we can only speculate that DNMT1 may compensate for the loss of *de novo* DNA methylation activities in establishing *Dnmt3a/3b-*DKO cEpiSCs. Considering the dual catalytic and noncatalytic function of DNMT1^51^, it is also an open question whether the failure to derive cEpiSCs from *Dnmt1/3a/3b-* TKO but not *Dnmt3a/3b-*DKO is due to the lack of DNA methylation dependent and/or independent functions.

## ACKNOWLEDGMENTS

We thank Dr. Taiping Chen for providing the DNMT KO ESCs, Dr. Guo-Liang Xu for providing the DNMT3A and DNMT3B antibodies for embryo staining, and Dr. Isao Suetake for providing the DNMT3L antibody for embryo staining. This work was supported by National Institutes of Health (NIH) to X.H (R21HD106263), J.W. (R01GM129157, R01HD095938, R01HD097268, and R01HL146664), and A.-K.H (R01DK127821, R01HD094868 and P30CA008748), and by contracts from New York State Stem Cell Science (NYSTEM) to J.W. (C35583GG and C35584GG).

## AUTHOR CONTRIBUTIONS

X.H. conceived, designed, and conducted the study, performed bioinformatics analysis, and wrote the manuscript; S.B. and A.-K.H. performed embryo staining; C.L. and H.W. performed mass spectrometry analysis; Y.X., Y.Z., and W.X. performed STEM-seq of mouse embryos and data analysis. V.M. and H.Z. provided reagents and contributed to experiments; J.W. conceived the project, designed the experiments, and prepared and approved the manuscript.

## STAR METHODS

### KEY RESOURCES TABLE

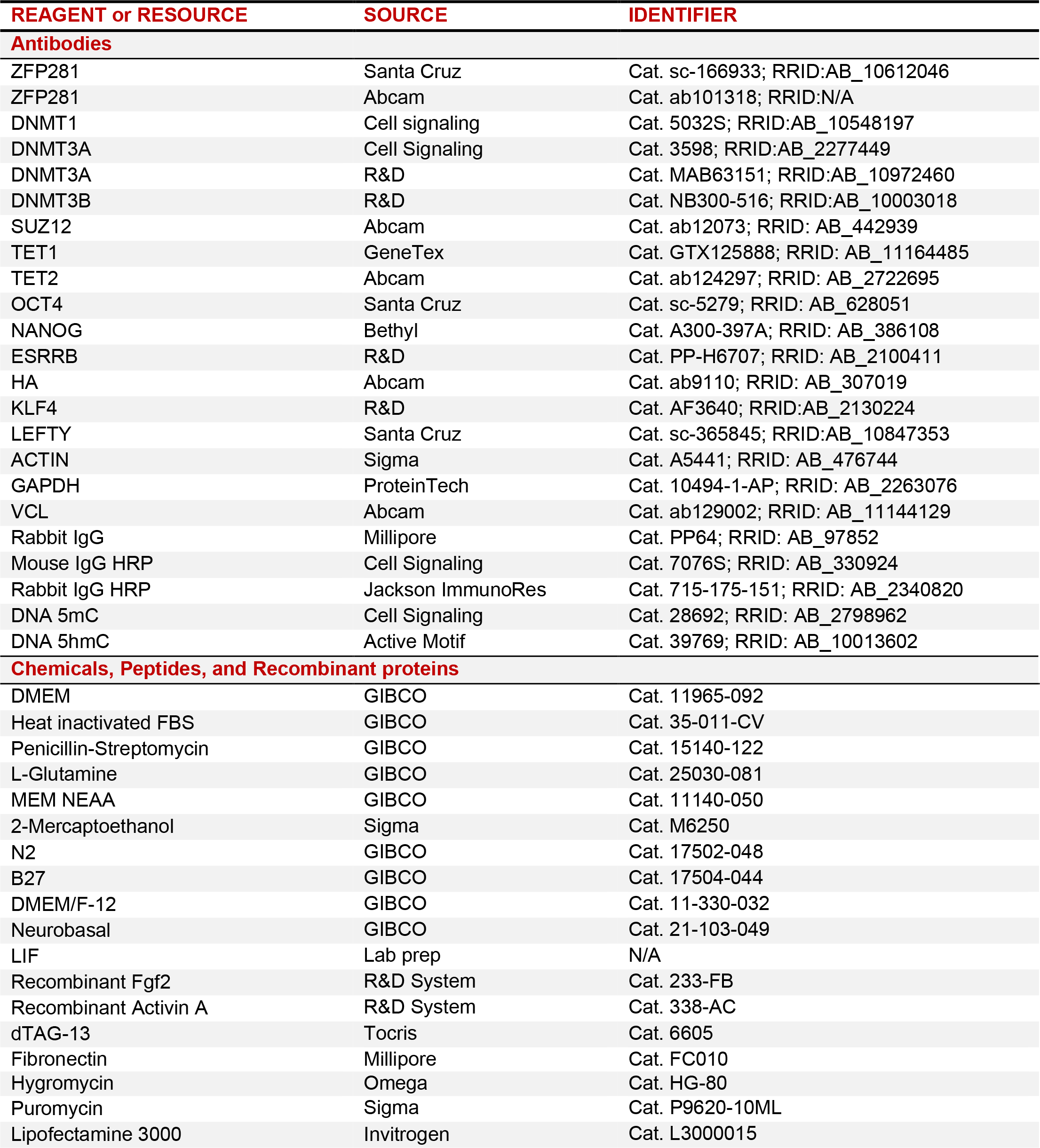
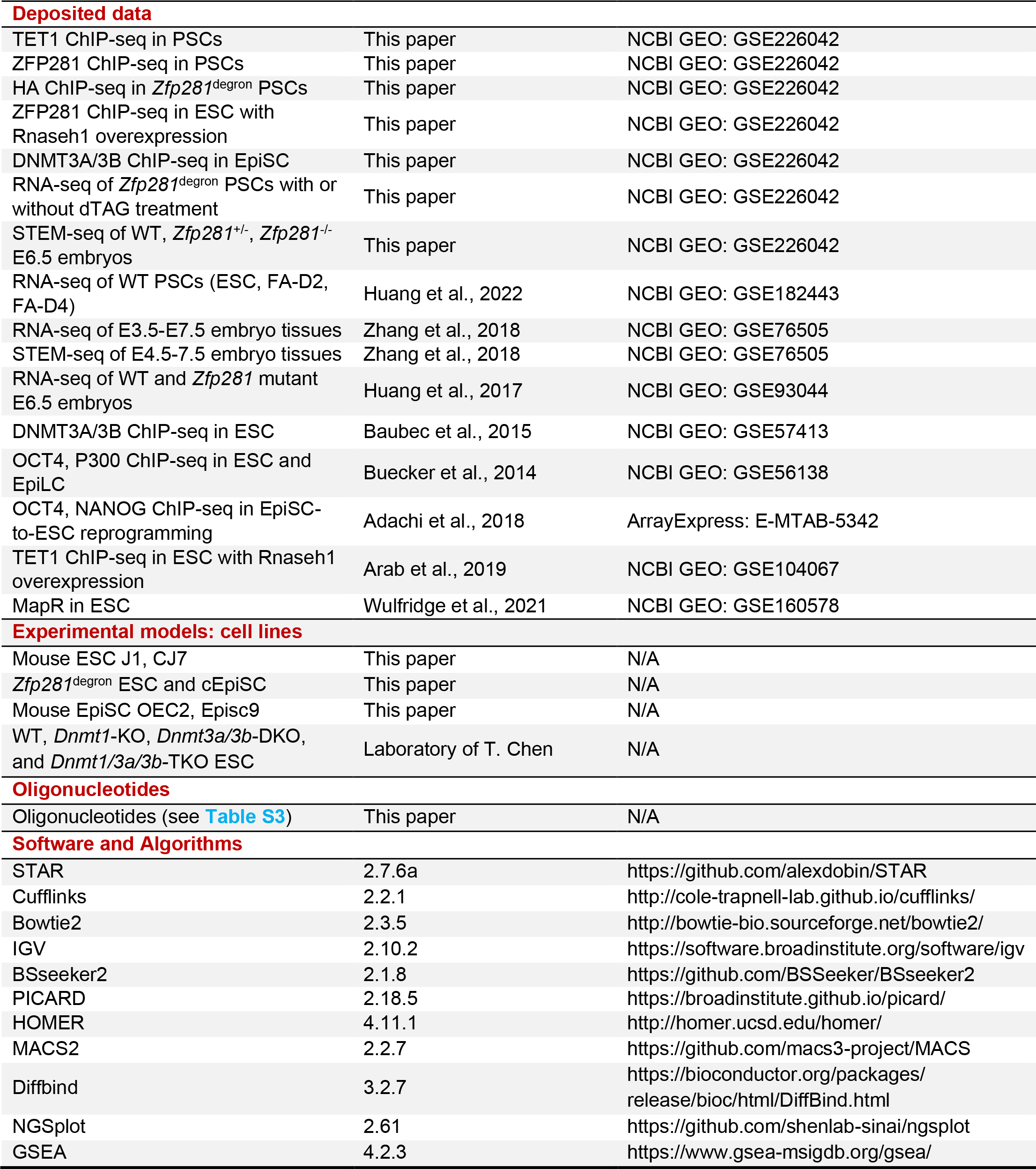

### RESOURCE AVAILABILITY

#### Lead contact

Further information and requests for resources and reagents should be directed to and will be fulfilled by the lead contact, Jianlong Wang.

#### Materials availability

The *Zfp281*^degron^ ESCs generated in this paper are available from the lead contact with a completed Materials Transfer Agreement.

#### Data and code availability

The ChIP-seq, RNA-seq, and STEM-seq data have been deposited at the Gene Expression Omnibus (GEO) with accession code: GSE226042. The deposited data are publicly available as of the date of publication. This paper analyzes existing, publicly available data. These accession numbers for the datasets are listed in the key resources table.

This paper does not report the original code.

Any additional information required to reanalyze the data reported in this paper is available from the lead contact upon request.

### EXPERIMENTAL MODEL AND SUBJECT DETAILS

#### Cell culture and pluripotent state transition

If not specified, mouse ESCs were cultured on 0.1% gelatin-coated plates and in ES medium: DMEM medium supplemented with 15% fetal bovine serum (FBS), leukemia inhibitory factor (LIF, lab prepared), 0.1 mM 2-mercaptoethanol, 2 mM L-glutamine, 0.1 mM MEM non-essential amino acids (NEAA), 1% nucleoside mix (100X stock), and 50 U/mL Penicillin/Streptomycin (P/S). Mouse EpiSCs were cultured on fibronectin-coated (10 μg/mL) plates and in serum-free N2B27 medium supplemented with 0.1 mM 2-mercaptoethanol, 2 mM L-glutamine, 0.1 mM NEAA, 50 U/mL P/S, and supplemented with fresh Fgf2 (12 ng/mL) and Activin A (20 ng/mL) (FA medium). ESCs were passed with 0.05% Trypsin/EDTA, and EpiSCs were passed with Accutase.

For ESC-EpiLC-EpiSC differentiation, ESCs were seeded on fibronectin-coated (10 μg/mL) plates in ES medium overnight and switched to FA medium the next day. The adapted cells in FA medium for 2 days (FA-D2) and 4 days (FA-D4) were collected. After more than two weeks of FA culture, adapted cells were considered converted EpiSCs (cEpiSCs).

#### Mouse study

The *Zfp281*KO mice^22^ were used in this study. All mice experiments were conducted in accordance with the guidelines approved by the Institutional Animal Care and Use Committee (IACUC) at Columbia University Irving Medical Center (PI: Jianlong Wang, protocol #AABD5612) and Memorial Sloan Kettering Cancer Center (PI: Hadjantonakis, protocol #03-12-017).

#### *Zfp281*^degron^ knock-in (KI) and protein degradation

The CRISPR/Cas9 system was used to engineer ESCs for protein degradation of ZFP281, as previously described.^52^ The 5’- and 3’-homology arms of *Zfp281* for C-terminal insertion were PCR amplified from genomic DNA. The 5’- and 3’-homology arms and FKBP12^F36V^-2xHA- mCherry fragment were assembled by Gibson Assembly 2x Master Mix (NEB, E2611S). The CRISPR gRNA was subcloned into the pSpCas9(BB)-2A-Puro (PX459) vector (gRNA sequence in Table S3). ESCs were transfected with the donor and CRISPR vectors using Lipofectamine 3000 (Invitrogen). After 2 days of puromycin selection, mCherry-positive cells were seeded on a 96-well plate with single-cell per well using the BD Influx Cell Sorter. Cells were expanded and genotyped by PCR. Two clones (#2 and #21) with a homozygous knock-in were further expanded and used for experiments. *Zfp281*^degron^ ESCs and cEpiSCs were treated with dTAG13 (500 nM in DMSO, Tocris, 6605) for degradation of ZFP281 protein.

### METHOD DETAILS

#### Single-cell quantitative immunofluorescence (qIF)

E6.5 embryos were collected, and qIF was performed as we previously described.^22^ Briefly, embryos were permeabilized in PBS-0.5% Triton for 20 min at room temperature, washed in PBS- 0.1% Triton (PBT), and blocked at 4°C o/n in PBT-3% BSA. Primary and secondary antibody staining was performed overnight at 4°C. Counterstaining with Hoechst and fluorophore-coupled phalloidin (Life Technologies, Carlsbad, CA) was performed for 1 hr at room temperature, and images were taken on a Zeiss LSM880 laser scanning confocal microscope. Fluorescence intensity levels were measured on data acquired with the same imaging parameters. Postimplantation nuclear protein levels were quantified using Imaris software (Bitplane) by manually creating individual nuclear surfaces for each cell and quantifying the fluorescence level inside the volume defined by these surfaces. Statistical significance was calculated on the average level of corrected fluorescence per embryo using an unpaired two-tailed Student T-test with Welch’s correction when standard deviations differed between samples. The following primary antibodies were used: ZFP281 (Santa Cruz, sc-166933), DNMT3A and DNMT3B (gift from Dr. Guo-Liang Xu), and DNMT3L (gift from Dr. Isao Suetake).

#### Small-scale TELP-enabled methylome sequencing (STEM-seq)

STEM-seq for low-input genome-wide DNA methylation profiling in epiblast cells of WT, *Zfp281*^+/-^, and *Zfp281*^-/-^ E6.5 embryos was performed as previously described.^1^ All STEM-seq datasets were mapped to the mm9 genome by BSSeeker2. Alignments were performed with the following parameters in addition to the default parameters: --bt2-p 8 --XS 0.2,3 --a CCCCCC --m 4. Multi-mapped reads and PCR duplicates were removed. After validating the reproducibility between replicates, we pooled data from replicates for subsequent analyses. For methylation analysis, the CG methylation was calculated as the total methylated counts (combining Watson and Crick strands) divided by the total counts across all reads covering that CG.

#### *Rnaseh1* overexpression

Mouse *Rnaseh1* coding sequence (CDS) was amplified from the cDNA of mouse ESCs by reverse transcription with SuperScript III Reverse Transcriptase (Invitrogen). Then *Rnaseh1* CDS was cloned into a piggyBac (PB) expression vector with Hygromycin. The PB-*Rnaseh1*-Hygro and empty PB-Hygro vectors were cotransfected with helper PBase (encodes the transposase) vector in ESCs with Lipofectamine 3000 (Invitrogen). Transfected ESCs were selected with 100 µg/ml Hygromycin for one week with at least two passages.

#### Western blot analysis

For Western blot analysis, total proteins were extracted by RIPA buffer with a protease inhibitor cocktail (Sigma). Protein concentrations were measured by Bradford assay (Pierce, 23236), balanced, and subjected to SDS-PAGE analysis. The following primary antibodies were used: ZFP281 (Santa Cruz, sc-166933), TET1 (GeneTex, GTX125888), DNMT1 (Cell signaling, 5032S), DNMT3A (R&D, MAB63151), DNMT3B (R&D, NB300-516), OCT4 (Santa Cruz, sc-5279), ESRRB (R&D, PP-H6707), KLF4 (R&D, AF3640), NANOG (Bethyl, A300-397A), TET2 (Abcam, ab124297), SUZ12 (Abcam, ab12073), LEFTY (Santa Cruz, sc-365845), ACTIN (Sigma, A5441), GAPDH (ProteinTech, 10494-1-AP), and Vinculin (VCL, Abcam, ab129002).

#### Dot blot analysis

The genomic DNA dot-blot analysis of 5mC and 5hmC was performed following the DNA Dot Blot Protocol (Cell Signaling, #28692), as previously described.^52^ Briefly, genomic DNA was extracted using Quick-DNA Miniprep Plus Kit (Zymo Research, D4068), and DNA concentration was measured by NanoDrop. The same amount of DNA was denatured with 10X DNA denaturing buffer (1 M NaOH and 0.1 M EDTA) and incubated at 95°C for 10 min, which was then immediately mixed with an equal volume of 20X SSC buffer, pH 7.0 (Invitrogen, 15557044) and chilled on ice. The DNA samples were diluted with a pre-determined amount and loaded on the positive-charged Nylon membrane (GE Amersham, RPN2020B) using a vacuum chamber (Manifold, SRC-96). The membrane was dried, auto-crosslinked with 1200 x100 μJ/cm^2^, and blocked with 5% milk/TBST for 1 h. Next, the membrane was incubated with 5mC (Cell Signaling, 28692) or 5hmC (Active Motif, 39769) antibodies, the same as the western blot analysis.

#### Genomic DNA 5mC and 5hmC quantification by mass spectrometry

The UHPLC-MS/MS analysis for 5mC and 5hmC quantification was performed as previously described^53^ on an Agilent 1290 Infinity II ultrahigh performance LC system coupled with an Agilent 6470 triple quadrupole mass spectrometer equipped with a jet stream electrospray ionization source (Santa Clara, CA). MS was operated under positive ionization using multiple reactions monitoring (MRM) mode: *m/z* 242->83 for 5mC and *m/z* 258->142 for 5hmC. The frequencies of 5mC and 5hmC over total deoxycytidine (dC) were calibrated by corresponding stable isotope-labeled internal standards.

#### Zfp281 shRNA knockdown

*Zfp281* knockdown in ESCs and EpiSCs was performed as previously described,^21^ with either a control empty vector or two independent *Zfp281* shRNAs.

#### RT-qPCR

Total RNA was extracted using the GeneJet RNA Purification Kit (Thermo Scientific, K0732). Reverse transcription was performed using the qScript kit (Quanta, 95048). Relative expression levels were determined using a QuantStudio 5 Real-Time PCR System (Applied Biosystems). Gene expression levels were normalized to *Gapdh*. Primers for RT-qPCR are listed in Table S3.

#### Chromatin immunoprecipitation (ChIP) and sequencing

ChIP was performed as previously described.^52^ Briefly, cell pellets were crosslinked with 1% (w/v) formaldehyde for 10 min at RT, followed by the addition of 125 mM glycine to stop the reaction.

Next, chromatin extracts were sonicated into 200–500 bp with Bioruptor Plus (settings of 30 sec ON, 30 sec OFF, 30 cycles) or with Bioruptor Pico (settings of 30 sec ON, 30 sec OFF, 15 cycles). ChIP was performed with the following primary antibodies: ZFP281 (Abcam, ab101318), HA (Abcam, ab9110), TET1 (GenTex, GTX125888), DNMT3A (Cell Signaling, 3598), DNMT3B (R&D, NB300-516), or rabbit IgG (Millipore, PP64) overnight at 4°C with continuous mixing, followed by incubation with protein G dynabeads (Invitrogen, 10004D) for another 2 hr at 4°C. The immunoprecipitated DNA was washed with ChIP RIPA buffer and purified with ChIP DNA Clean & Concentrator columns (Zymo Research, D5205). ChIP-qPCR was performed with Roche SYBR Green reagents and a LightCycler480 (Roche) machine. Percentages of input recovery were calculated. The ChIP-qPCR primers are listed in Table S3.

For ChIP-seq, 10% of sonicated genomic DNA was used as ChIP input. Libraries were prepared using the NEBNext Ultra II DNA library prep kit and index primers sets (NEB, 7645S, E7335S) following the standard protocol. Sequencing was performed with the Illumina HiSeq 4000 Sequencer according to the manufacturer’s protocol. Libraries were sequenced as 150-bp paired-end reads.

### RNA sequencing

Total RNA was extracted using the GeneJet RNA Purification Kit (Thermo Scientific, K0732). RNA-seq library construction was performed at Novogene with a standard polyA-enrichment protocol. Sequencing was performed on an Illumina HiSeq 4000 Sequencer, and 150-bp paired-end reads were obtained.

### QUANTIFICATION AND STATISTICAL ANALYSIS

#### ChIP-seq data processing

All ChIP-seq reads were pre-processed by trim_galore (v0.6.3) and aligned to the mm9 mouse genome using the bowtie2 (v2.3.4) program, and the parameters were “-X 1000 --no-mixed --no-discordant”. The aligned reads were exported (-F 0x04 -f 0x02) and sorted with samtools. Duplicates were removed with MarkDuplicates function in the PICARD (v2.14.0) package. The aligned ChIP-seq bam files of ZFP281 ChIP in WT PSCs and HA-ChIP in *Zfp281*^degron^ PSCs were combined. All bam files were converted to a binary tiled file (tdf) and visualized using IGV (v2.7.2) software.

ChIP-seq peaks were determined by the MACS2 program (v.2.2.7), using the input ChIP-seq as the control data, and all other parameters were the default settings. ChIP-seq peaks were annotated using the annotatePeaks module in the HOMER program (v4.11) against the mm9 genome. Motif analysis was performed using the findMotifsGenome module in HOMER against the mm9 genome, and with parameters: -size given -len 10. A target gene of a called peak was defined as the nearest gene’s transcription start site (TSS) with a distance to TSS less than 5 kb. Heatmaps and mean intensity curves of ChIP-seq data at specific genomic regions were plotted by the NGSplot program (v2.61) centered by the middle point “(start+end)/2” of each region. Diffbind (v3.2.7) was used to compare the intensity of reads at specific regions between different ChIP-seq data. Peaks with significantly different intensities were determined by FDR < 0.05.

#### RNA-seq data processing

For RNA-seq data processing, reads were aligned to the mouse genome mm9 using STAR (v2.7.6a) with the default settings. Transcript assembly and differential expression analyses were performed using Cufflinks (v2.2.1). Assembly of novel transcripts was not allowed (-G). Other parameters of Cufflinks were the default setting. The summed FPKM (fragments per kilobase per million mapped reads) of transcripts sharing each gene_id was calculated and exported by the Cuffdiff program. Differentially expressed genes (DEGs) were determined by two-sided T-test P-value<0.05 and fold-change>2 or by Q-value (FDR) < 0.05. Volcano plots for gene expression by fold change versus P-value were generated using R.

For hierarchical clustering analysis, the gene expression table was imported by Cluster 3.0 software. Mean-center and normalization of gene expression were performed, then the analysis was performed with average-linkage of genes. Clustered genes with normalized expression values (z-score) were shown in Heatmap with the Java TreeView (v1.1.6) program.

For principal component analysis (PCA), batch effects were adjusted by the *ComBat* function implemented in the *sva* Bioconductor package (v.3.18.0). PCA was performed with the Cluster 3.0 software. PC values were visualized with the plot3d function in the rgl package using R (v4.1.0) scripts.

#### Gene set enrichment analysis (GSEA) and gene ontology (GO) analysis

GSEA (v4.2.3) was used to determine the statistically enriched gene sets by comparing the WT and untreated *Zfp281*^degron^ EpiSCs (Ctrl: 4 samples) and the 2 days and 4 days of dTAG-treated *Zfp281*^degron^ EpiSCs (dTAG: 4 samples). The curated C5: ontology gene sets were downloaded from https://www.gsea-msigdb.org/gsea/. GSEA enrichment plot, normalized enrichment score (NES), and Q-value (FDR) were indicated for each enrichment test.

Gene ontology (GO) analyses were performed using the DAVID gene ontology functional annotation tool (https://david.ncifcrf.gov/tools.jsp) with all *Mus musculus* genes as a reference list.

#### Single-cell analysis of mouse embryo tissues

The scRNA-seq data^43^ with detailed annotation of gastrulating mouse embryo tissues (E6.5-E8.5) were available at https://marionilab.cruk.cam.ac.uk/MouseGastrulation2018/. Expression maps of gene-of-interest were downloaded by projection type: UMAP; cell subset: all timepoints; plot color: cell type.

#### Statistical analysis

If not specified, qPCR analysis was performed in technical triplicates, and the error bars indicate the standard deviation of the mean. The p-values were calculated using an unpaired two-sided T-test using Excel software. The line plots in Figure 1C-D represent the mean expression value of each cluster with a 95% confidence interval (CI) using the GraphPad Prism software (v9.2.0). The boxplots in Figures 3F and 6F present the 25th, median, and 75th quartiles, and the whiskers extend 1.5 of interquartile ranges. The p-value was calculated from the two-sided Mann-Whitney test using R. The scatter plots in Figure S6B calculated the log2 ratio of gene expression (D4/D0 vs. D2/D0), and a linear regression line and coefficient of determination (R^2^) value were calculated by Excel software. If not specified, statistical analysis was performed with R (v4.1.0) scripts on the R-Studio platform (v1.4.1). The statistical details of the experiment are indicated in the figure legend.

## Notes

### Competing Interest Statement

The authors have declared no competing interest.

