## Supplementary material for "ZFP281 coordinates DNMT3 and TET1 for transcriptional and epigenetic control in pluripotent state transitions": SI Figures and Legends

### Figure S1

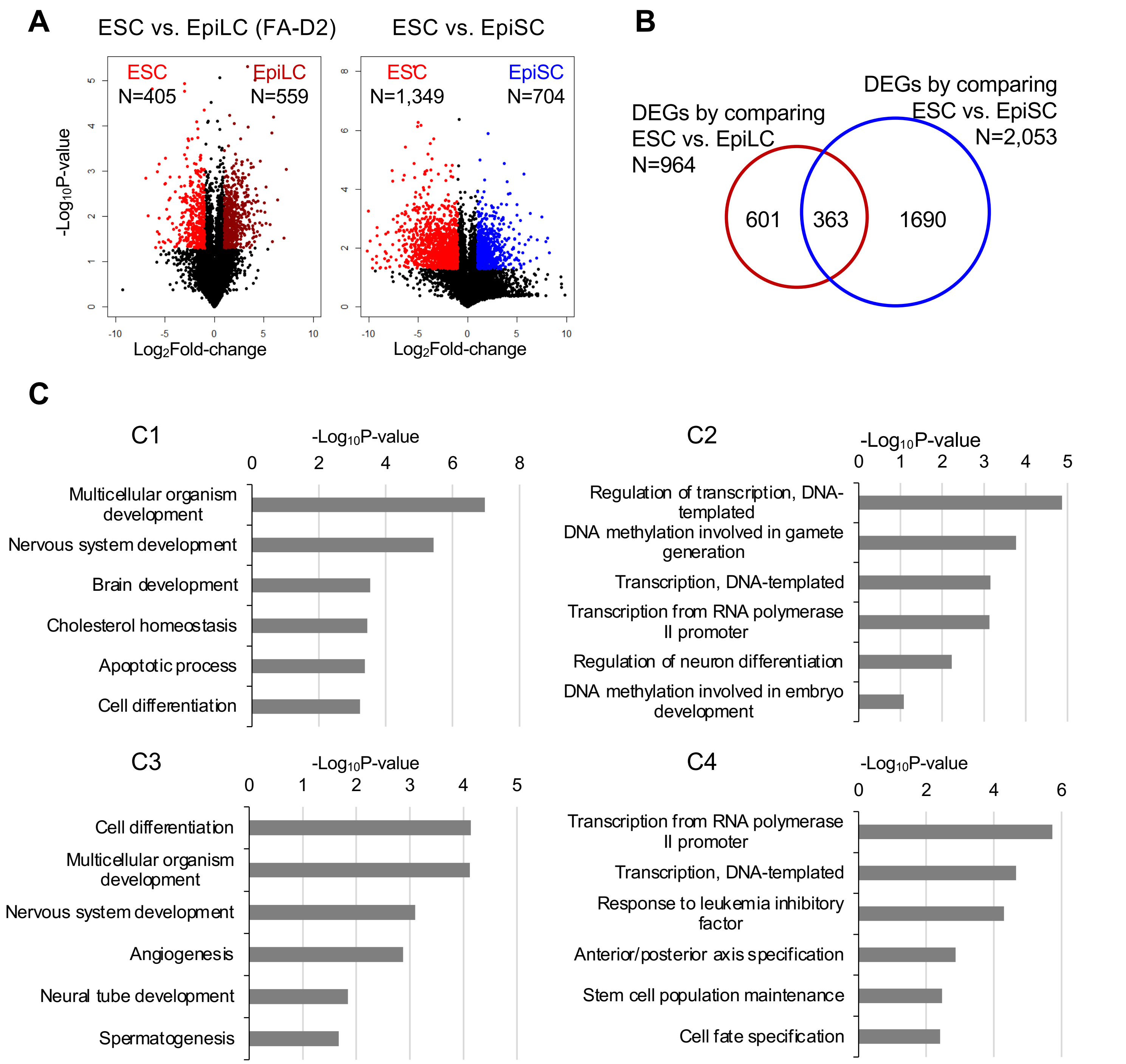
A
ESC vs. EpiLC (FA-D2)
ESC vs. EpiSC
ESC
N=405
EpiLC
N=559
ESC
N=1,349
EpiSC
N=704
-Log10P-value
Log2Fold-change
B
DEGs by comparing ESC vs. EpiLC N=964
DEGs by comparing ESC vs. EpiSC N=2,053
601
363
1690
C
C1
-Log10P-value
Multicellular organism development
Nervous system development
Brain development
Cholesterol homeostasis
Apoptotic process
Cell differentiation
C2
-Log10P-value
Regulation of transcription, DNA-templated
DNA methylation involved in gamete generation
Transcription, DNA-templated
Transcription from RNA polymerase II promoter
Regulation of neuron differentiation
DNA methylation involved in embryo development
C3
-Log10P-value
Cell differentiation
Multicellular organism development
Nervous system development
Angiogenesis
Neural tube development
Spermatogenesis
C4
-Log10P-value
Transcription from RNA polymerase II promoter
Transcription, DNA-templated
Response to leukemia inhibitory factor
Anterior/posterior axis specification
Stem cell population maintenance
Cell fate specification

#### SUPPLEMENTARY FIGURE LEGENDS

**Figure S1. Dynamic gene expression during the pluripotent state transitions. Related to Figure 1.**

(A) Volcano plots depicting DEGs (P-value<0.05, fold-change>2) by comparing the ESC vs. EpiLC (FA-D2, left) and ESC vs. EpiSC (right) samples. Numbers of the DEGs highly expressed in ESCs or EpiLCs/EpiSCs were indicated.

(B) Ven diagram depicting the overlapped genes between DEGs by comparing ESC vs. EpiLC and ESC vs. EpiSC samples.

(C) Gene ontology (GO) analysis for the C1~C4 genes identified from Figure 1B.

Figure S2

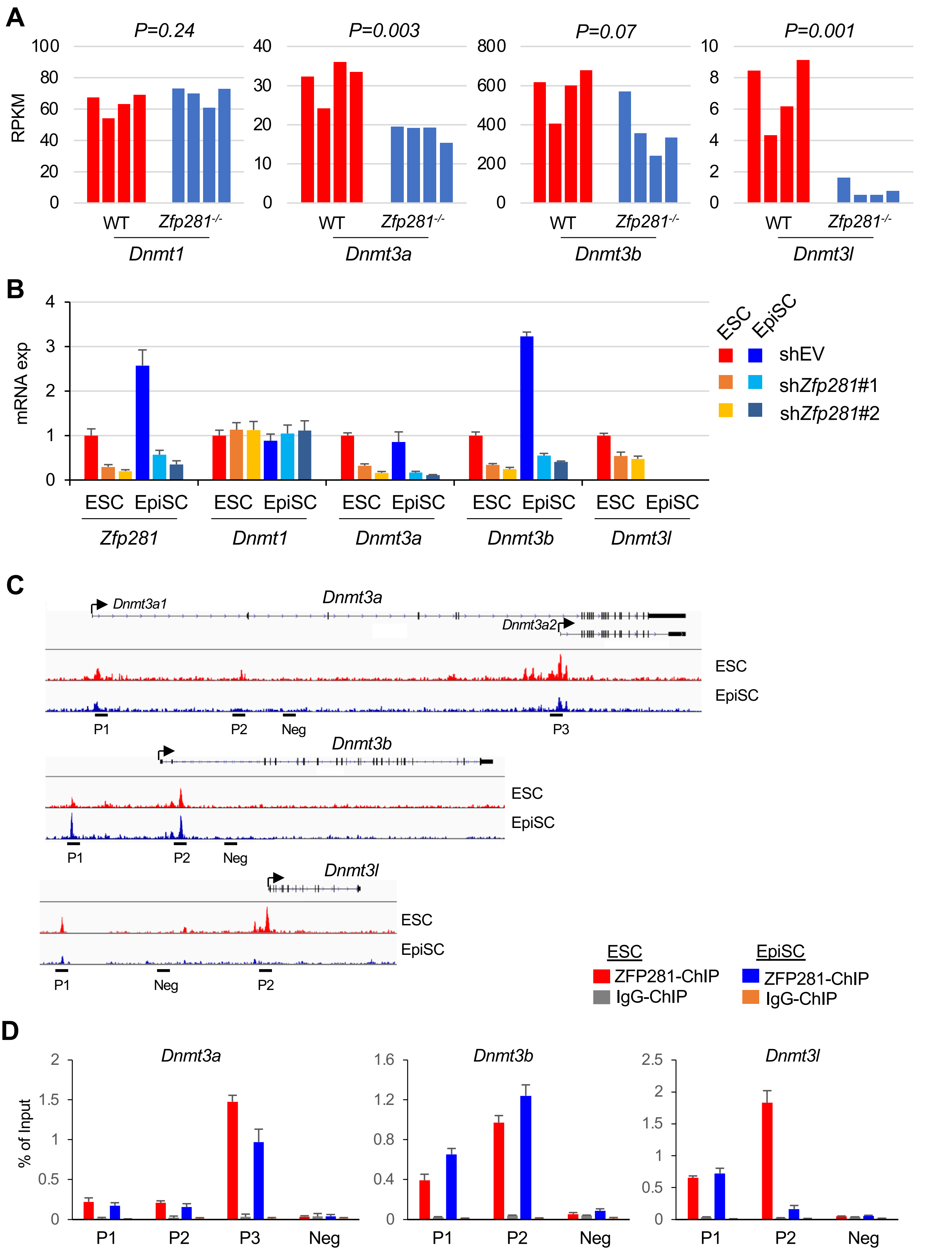
A
P=0.24
P=0.003
P=0.07
P=0.001
RPKM
WT
Zfp281-/-
Dnmt1
Dnmt3a
Dnmt3b
Dnmt3l
B
mRNA exp
ESC
EpiSC
shEV
shZfp281#1
shZfp281#2
Zfp281
Dnmt1
Dnmt3a
Dnmt3b
Dnmt3l
C
Dnmt3a1
Dnmt3a
Dnmt3a2
ESC
EpiSC
P1
P2
Neg
P3
Dnmt3b
Dnmt3l
ESC
ZFP281-ChIP
IgG-ChIP
EpiSC
ZFP281-ChIP
IgG-ChIP
D
% of Input
Dnmt3a
Dnmt3b
Dnmt3l
P1
P2
P3
Neg

**Figure S2. ZFP281 transcriptionally regulates Dnmt3a/3b expression. Related to Figure 2.**

(A) Expression of DNMT family genes in WT and *Zfp281*$^{-/-}$ E6.5 embryos by RNA-seq analysis (from our published study[22]).

(B) Expression of DNMT family genes in ESCs and EpiSCs upon *Zfp281* knockdown by shRNAs. Empty vector (EV) and two independent *Zfp281* shRNAs were used for the knockdown.

(C-D) ChIP-seq tracks (C) and ChIP-qPCR (D) analysis depicting binding of ZFP281 at *Dnmt3a*, *Dnmt3b*, and *Dnmt3l* regulatory loci. The *Dnmt3a* gene has two TSSs and transcribes *Dnmt3a1* (long) and *Dnmt3a2* (short) isoforms. ChIP-qPCR was performed targeting the positive (P1, P2, P3) and negative (Neg) sites.

Figure S3

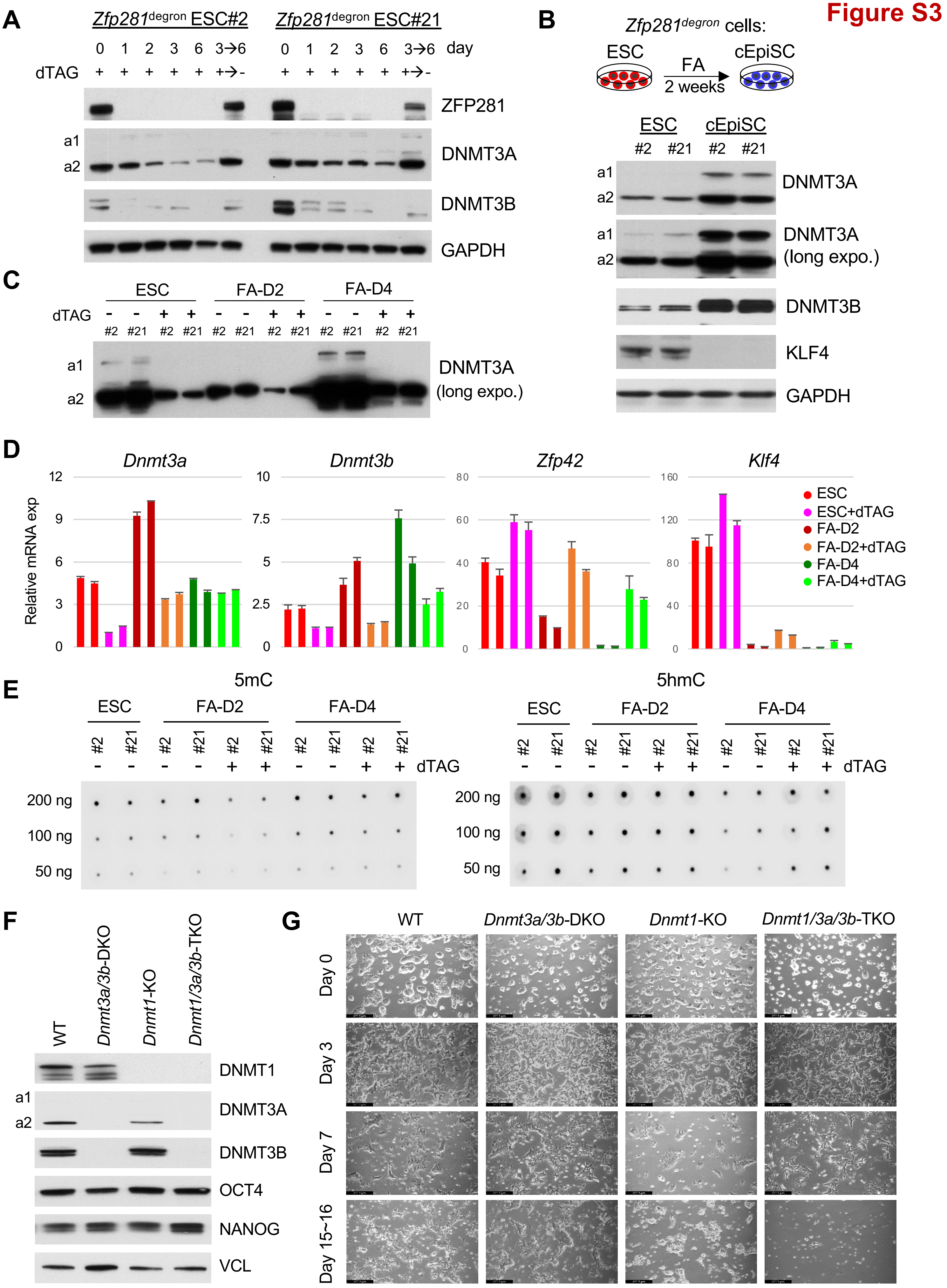

A
Zfp281degron ESC#2
Zfp281degron ESC#21
0 1 2 3 6 3→6 day
dTAG + + + + + +→-
ZFP281
a1
a2
DNMT3A
DNMT3B
GAPDH
B
Zfp281degron cells:
ESC
FA
2 weeks
cEpiSC
ESC
cEpiSC
#2 #21 #2 #21
DNMT3A
DNMT3A (long expo.)
DNMT3B
KLF4
GAPDH
C
ESC
FA-D2
FA-D4
dTAG
#2 #21
DNMT3A (long expo.)
D
Dnmt3a
Dnmt3b
Zfp42
Klf4
Relative mRNA exp
ESC
ESC+dTAG
FA-D2
FA-D2+dTAG
FA-D4
FA-D4+dTAG
E
5mC
5hmC
200 ng
100 ng
50 ng
dTAG
F
WT
Dnmt3a/3b-DKO
Dnmt1-KO
Dnmt1/3a/3b-TKO
DNMT1
DNMT3A
DNMT3B
OCT4
NANOG
VCL
G
Day 0
Day 3
Day 7
Day 15~16

**Figure S3. ZFP281 activates Dnmt3a/3b expression and DNA methylation in pluripotent state transitions. Related to Figure 3.**

(A) ZFP281 depletion reduced DNMT3A and DNMT3B proteins in ESCs. *Zfp281*$^{degron}$ ESCs were treated with dTAG continuously for 6 days (day 0, 1, 2, 3, 6 samples were shown) or for the first 3 days followed by 3 days without dTAG (3→6 days; +→- dTAG; day 6 sample was shown). Total lysates from the days indicated were prepared for western blot analyses.

(B) Western blots of DNMT3A and DNMT3B in *Zfp281*$^{degron}$ ESCs and cEpiSCs. Two isoforms of DNMT3A were labeled at different exposure times, and KLF4 was used as ESC-specific control.

(C) Western blot of DNMT3A with both isoforms under long exposure (related to Figure 3C).

(D) RT-qPCR analysis of *Dnmt3a, Dnmt3b*, and naive pluripotency markers *Klf4* and *Zfp42* in *Zfp281*$^{degron}$ PSCs with or without dTAG treatment.

(E) DNA dot-blot analysis for 5mC (left) and 5hmC (right) intensities in the genomic DNA of *Zfp281*$^{degron}$ PSCs with or without dTAG treatment.

(F) Western blots of DNMT1/3A/3B in WT, *Dnmt3a/3b*-DKO, *Dnmt1*-KO, and *Dnmt1/3a/3b*-TKO ESCs. Expression of OCT4 and NANOG indicated these DNMT KO ESCs maintain pluripotency. VCL (Vinculin) served as a loading control.

(G) Images of WT, *Dnmt3a/3b*-DKO, *Dnmt1*-KO, and *Dnmt1/3a/3b*-TKO ESCs differentiating to EpiSCs at different time points. At Day 15~16, only *Dnmt1/3a/3b*-TKO cells cannot be maintained in primed cultures. Scale bar: 277 μm.

Figure S4

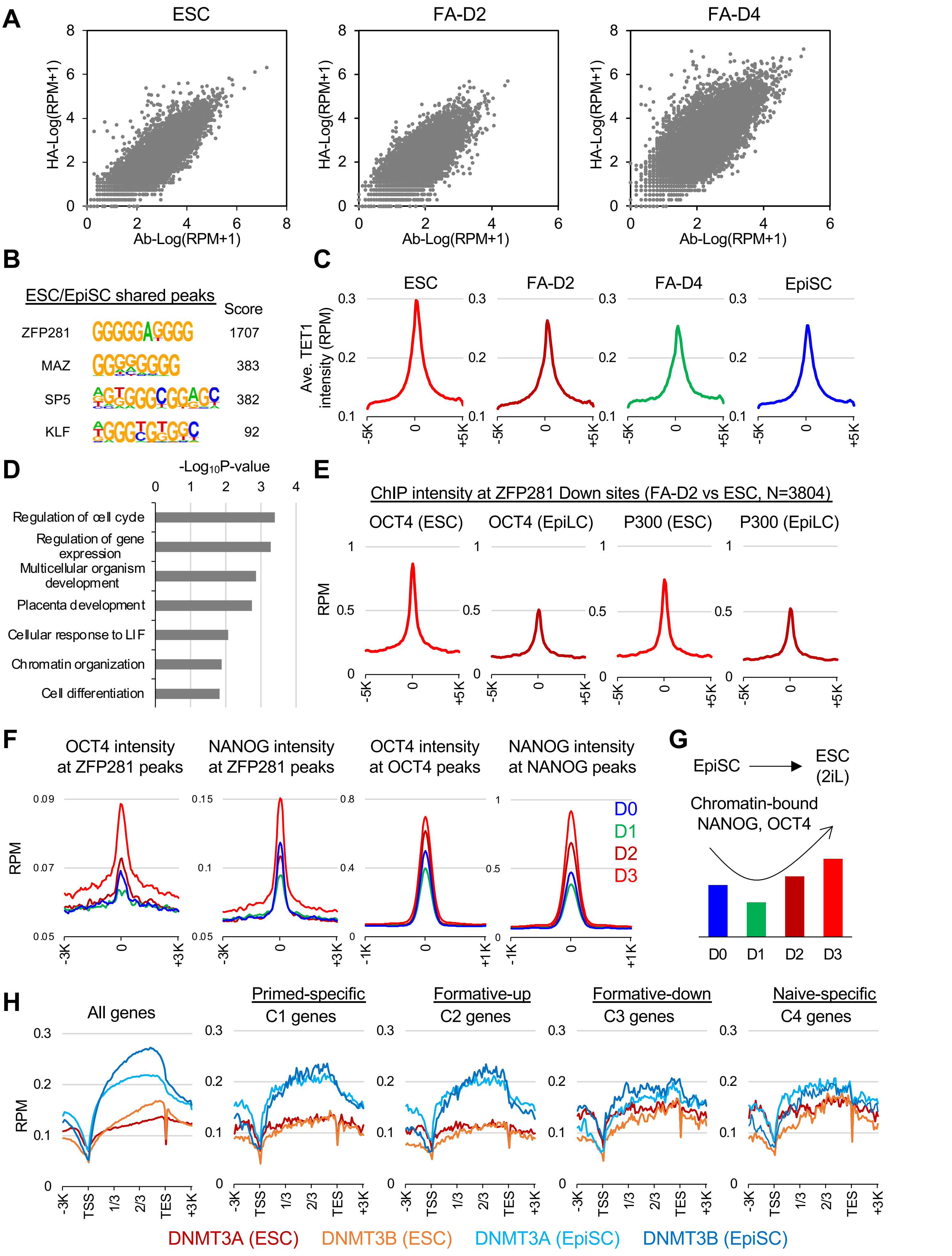
A
ESC
FA-D2
FA-D4
HA-Log(RPM+1)
Ab-Log(RPM+1)
B
ESC/EpiSC shared peaks
Score
ZFP281 1707
MAZ 383
SP5 382
KLF 92
C
Ave. TET1 intensity (RPM)
ESC
FA-D2
FA-D4
EpiSC
D
-Log10P-value
Regulation of cell cycle
Regulation of gene expression
Multicellular organism development
Placenta development
Cellular response to LIF
Chromatin organization
Cell differentiation
E
ChIP intensity at ZFP281 Down sites (FA-D2 vs ESC, N=3804)
OCT4 (ESC)
OCT4 (EpiLC)
P300 (ESC)
P300 (EpiLC)
RPM
F
OCT4 intensity at ZFP281 peaks
NANOG intensity at ZFP281 peaks
OCT4 intensity at OCT4 peaks
NANOG intensity at NANOG peaks
RPM
D0
D1
D2
D3
G
EpiSC
ESC (2iL)
Chromatin-bound NANOG, OCT4
D0 D1 D2 D3
H
All genes
Primed-specific C1 genes
Formative-up C2 genes
Formative-down C3 genes
Naive-specific C4 genes
RPM
-3K TSS 1/3 2/3 TES +3K
DNMT3A (ESC)
DNMT3B (ESC)
DNMT3A (EpiSC)
DNMT3B (EpiSC)

**Figure S4. Dynamic chromatin occupancy of ZFP281/TET1 and feedback transcriptional control of DNMT3A/3B in pluripotent state transitions. Related to Figure 4.**

(A) Scatter plots depicting the correlation of ZFP281 peak intensities by $\log_2$(RPM+1) from HA-ChIP and ZFP281 antibody (Ab)-ChIP experiments in ESC, FA-D2, and FA-D4 samples.

(B) The top enriched motifs identified from the ESC/EpiSC-shared ZFP281 peaks.

(C) Mean intensity plots depicting TET1 ChIP-seq intensity at all TSSs (within ± 5K bp) in different pluripotent states.

(D) Gene ontology (GO) analysis for the target genes from the 945 bimodal ZFP281 peaks (TSS < 5K bp).

(E) OCT4 (left) and P300 (right) ChIP intensities at ZFP281 peaks with decreased intensity in FA-D2 EpiLCs compared with those in ESCs (N=3804, from Figure 4F, left panel). OCT4 and P300 ChIP-seq data were from a published study.[20]

(F) OCT4 and NANOG ChIP intensities at ZFP281 peak regions (within ± 3K bp around peak center) and OCT4 or NANOG peak regions (within ± 1K bp around peak center). OCT4 and NANOG ChIP-seq data were from a published study.[32]

(G) Schematic depiction of the chromatin-bound NANOG and OCT4 during the 3-day EpiSC-to-ESC reprogramming process.

(H) Mean intensity plots depicting DNMT3A and DNMT3B ChIP-seq intensities across all gene bodies (from the transcription start site/TSS to transcription termination site/TES, and extended 3K bp before TSS and after TES) in ESCs and EpiSCs. ChIP intensities were plotted at all genes and C1~C4 genes. DNMT3A/3B ChIP-seq data in ESCs were from a published study.[33]

### Figure S5

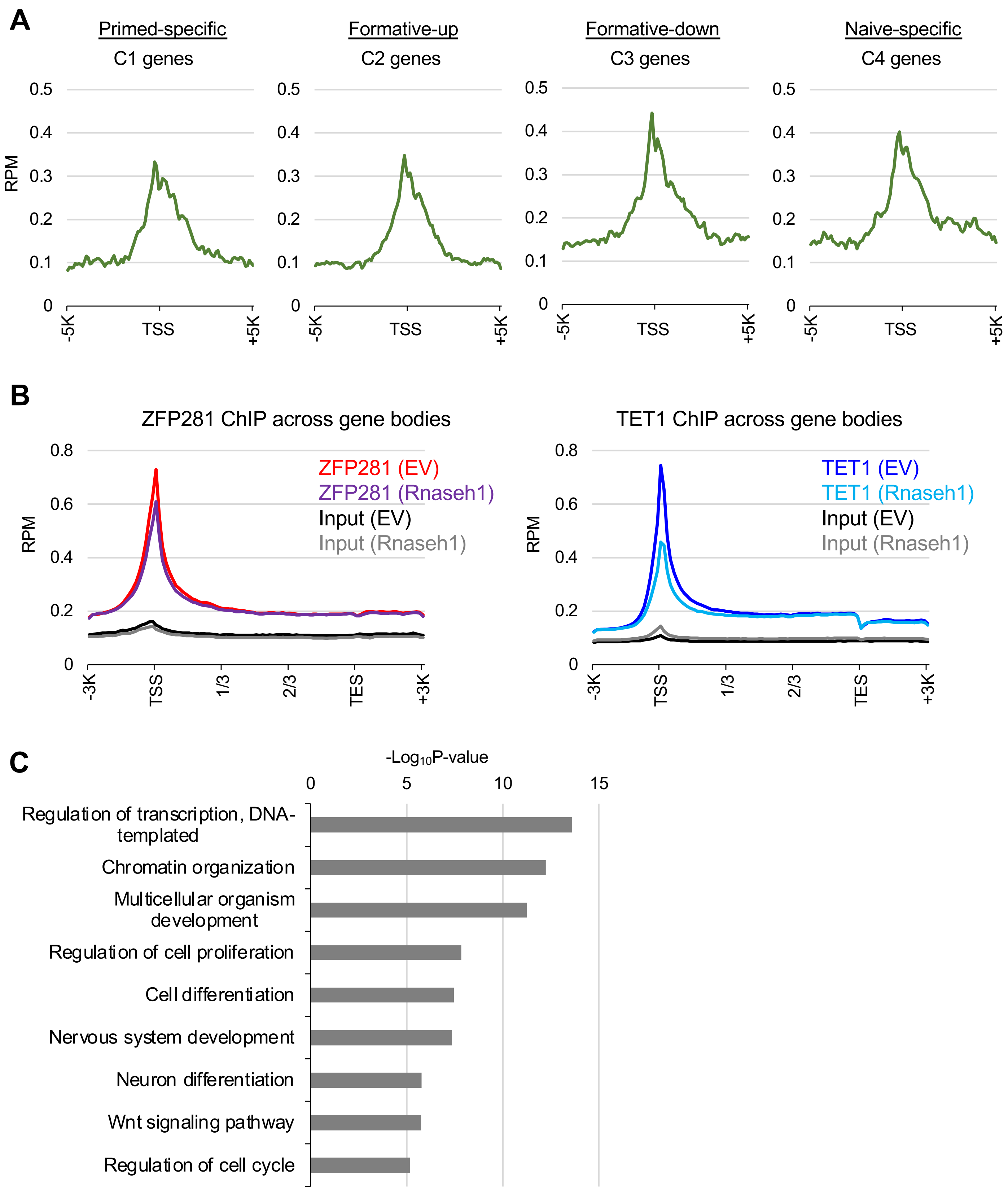
A
Primed-specific
C1 genes
Formative-up
C2 genes
Formative-down
C3 genes
Naive-specific
C4 genes
RPM
0.5
0.4
0.3
0.2
0.1
0
-5K
TSS
+5K
B
ZFP281 ChIP across gene bodies
ZFP281 (EV)
ZFP281 (Rnaseh1)
Input (EV)
Input (Rnaseh1)
TET1 ChIP across gene bodies
TET1 (EV)
TET1 (Rnaseh1)
Input (EV)
Input (Rnaseh1)
RPM
0.8
0.6
0.4
0.2
0
-3K
TSS
1/3
2/3
TES
+3K
C
-Log10P-value
0
5
10
15
Regulation of transcription, DNA-templated
Chromatin organization
Multicellular organism development
Regulation of cell proliferation
Cell differentiation
Nervous system development
Neuron differentiation
Wnt signaling pathway
Regulation of cell cycle

**Figure S5. ZFP281 chromatin association depends on the formation of R-loops. Related to Figure 5.**

(A) Mean intensity plots depicting the R-loop intensity by MapR at TSSs (within ± 5K bp) of C1~C4 genes in ESCs. MapR data in ESCs were from a published study.[37]

(B) ZFP281 and TET1 ChIP-seq intensities across all gene bodies (from transcription start site/TSS to transcription termination site/TES and extended 3K bp before TSS and after TES). ChIP input and ZFP281/TET1 ChIP intensities upon empty vector (EV) transfection or *Rnaseh1* overexpression were plotted. TET1 ChIP-seq data were from a published study.[38]

(C) Gene ontology (GO) analysis for the R-loop-sensitive ZFP281 target genes (TSS < 5K bp).

#### Figure S6

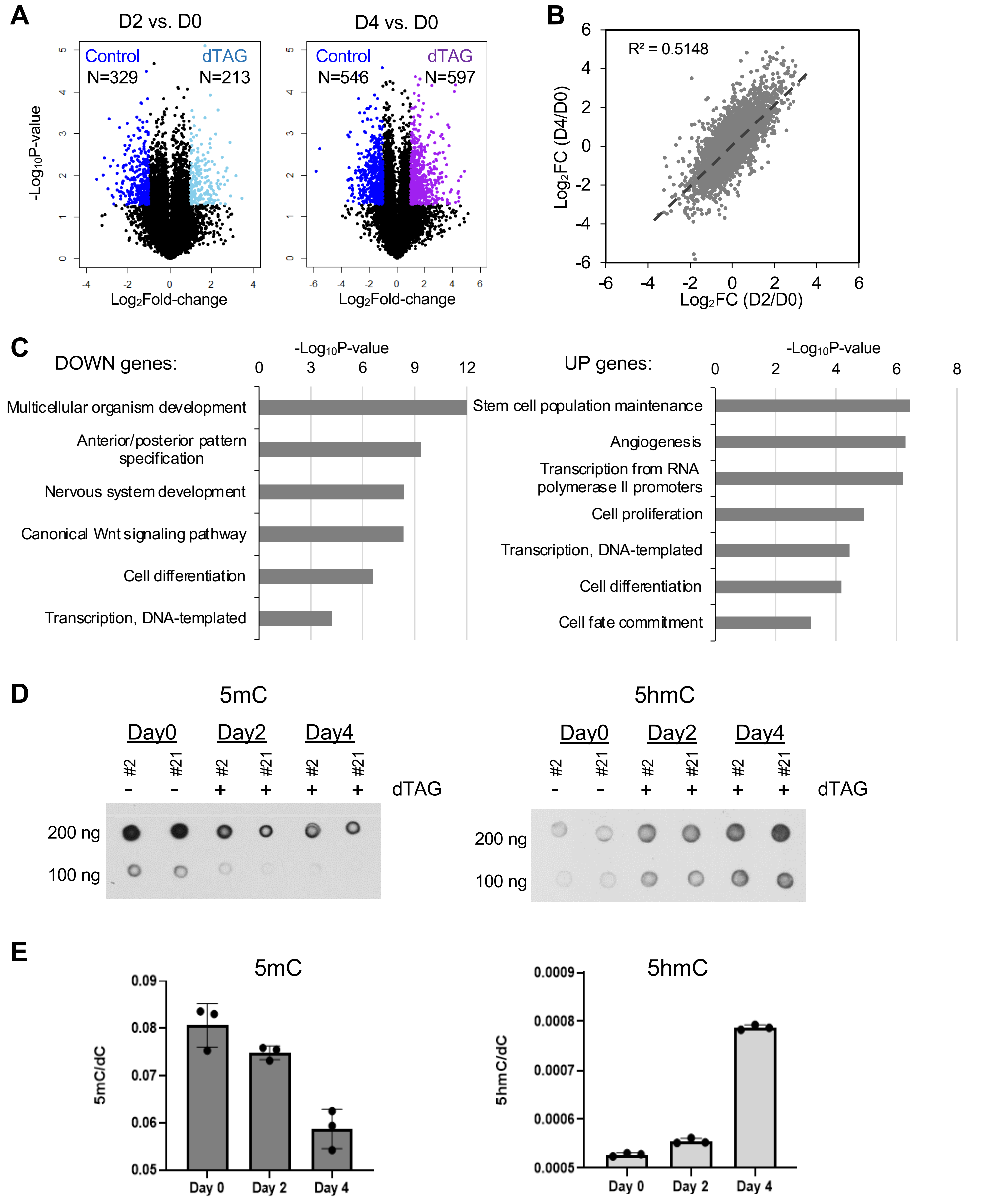
A
D2 vs. D0
D4 vs. D0
Control N=329
dTAG N=213
Control N=546
dTAG N=597
-Log10P-value
Log2Fold-change
B
R² = 0.5148
Log2FC (D4/D0)
Log2FC (D2/D0)
C
DOWN genes:
-Log10P-value
Multicellular organism development
Anterior/posterior pattern specification
Nervous system development
Canonical Wnt signaling pathway
Cell differentiation
Transcription, DNA-templated
UP genes:
-Log10P-value
Stem cell population maintenance
Angiogenesis
Transcription from RNA polymerase II promoters
Cell proliferation
Transcription, DNA-templated
Cell differentiation
Cell fate commitment
D
5mC
5hmC
Day0
Day2
Day4
#2
#21
dTAG
200 ng
100 ng
E
5mC
5hmC
5mC/dC
5hmC/dC
Day 0
Day 2
Day 4

**Figure S6. ZFP281 safeguards the homeostasis of DNA methylation and demethylation in maintaining primed pluripotency. Related to Figure 6.**

(A) Volcano plots depicting DEGs (P-value<0.05, fold-change>2) by comparing the *Zfp281*$^{degron}$ cEpiSCs treated with dTAG for 2 days (D2, left) and 4 days (D4, right) vs. the untreated (D0) cells. Numbers of the DEGs highly expressed in control or dTAG-treated cEpiSCs were indicated.

(B) Scatter plot depicting the relative gene expression upon 2 days (D2/D0) and 4 days (D4/D0) of dTAG treatment vs. D0 before the treatment in *Zfp281*$^{degron}$ cEpiSCs. A linear regression line and coefficient of determination ($R^2$) value were indicated.

(C) Gene ontology (GO) analysis for the Down- (left) and Up- (right) regulated genes (identified from Figure 6A) in *Zfp281*$^{degron}$ cEpiSCs with dTAG treatment.

(D-E) DNA dot-blot analysis (D) and UHPLC-MS/MS quantification (E) for 5mC and 5hmC in the genomic DNA of *Zfp281*$^{degron}$ cEpiSCs (two clones, #2 and #21) with dTAG treatments. In MS quantification, the intensity of 5mC or 5hmC over deoxycytidine (dC) was measured. Experiments were performed with technical triplicates.

**Figure S7**

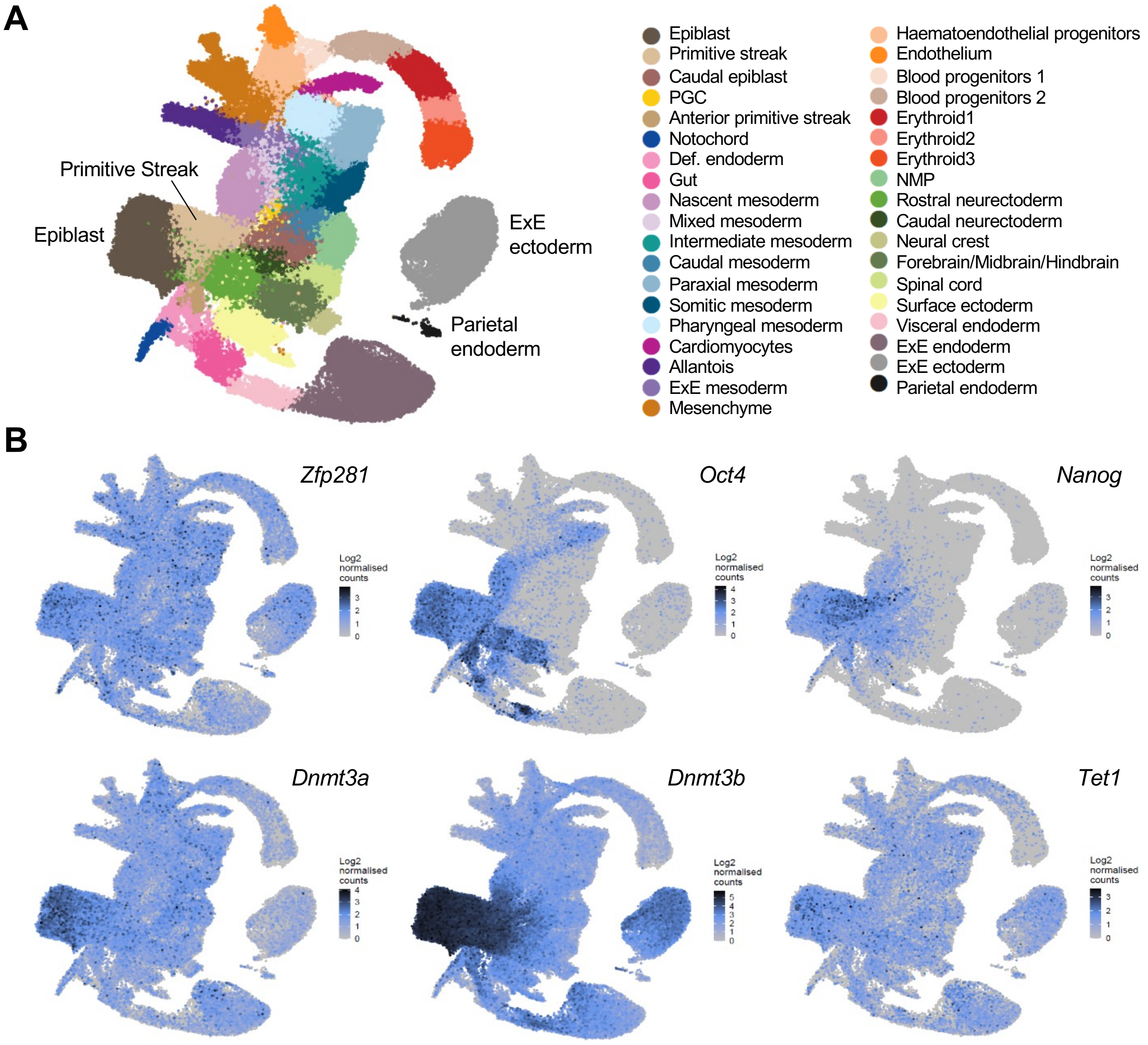
A
Primitive Streak
Epiblast
ExE
ectoderm
Parietal
endoderm
Epiblast
Primitive streak
Caudal epiblast
PGC
Anterior primitive streak
Notochord
Def. endoderm
Gut
Nascent mesoderm
Mixed mesoderm
Intermediate mesoderm
Caudal mesoderm
Paraxial mesoderm
Somitic mesoderm
Pharyngeal mesoderm
Cardiomyocytes
Allantois
ExE mesoderm
Mesenchyme
Haematoendothelial progenitors
Endothelium
Blood progenitors 1
Blood progenitors 2
Erythroid1
Erythroid2
Erythroid3
NMP
Rostral neurectoderm
Caudal neurectoderm
Neural crest
Forebrain/Midbrain/Hindbrain
Spinal cord
Surface ectoderm
Visceral endoderm
ExE endoderm
ExE ectoderm
Parietal endoderm
B
Zfp281
Oct4
Nanog
Dnmt3a
Dnmt3b
Tet1
Log2 normalised counts

**Figure S7. *Zfp281*, but not *Oct4* or *Nanog*, is broadly expressed together with *Dnmt3a/3b* and *Tet1* from E6.5 epiblast towards different lineage-specific precursors. Related to Figure 7.**

(A) Overall UMAP distribution (left) and annotation (right) of scRNA-seq (from a published study[43]) analysis of mouse embryos during gastrulation (E6.5-8.5). The E6.5 lineages (epiblast, primitive streak, ExE ectoderm, and parietal endoderm) are labeled in the UMAP.

(B) Expression of *Zfp281, Oct4, Nanog, Dnmt3a, Dnmt3b*, and *Tet1* from the scRNA-seq data. *Zfp281, Dnmt3a/b*, and *Tet1* are broadly expressed in E6.5 pluripotent epiblast and different lineage-specific precursors.
